# An emergent biosynthetic pathway to essential amino acids by metabolic metathesis

**DOI:** 10.1101/2023.09.06.556532

**Authors:** Julie Rivollier, Sandrine Gosling, Valérie Pezo, Marie-Pierre Heck, Philippe Marlière

**Author notes:** Altar SAS, 91000 Evry, France. CEA, Genoscope, 91000 Evry, France.

## Abstract

An experimental approach to implanting foreign chemical reactions into living cells is to select for the catalytic production of a vital building-block such as an amino acid. Alkene metathesis is unknown in extant biochemistry, but is emerging as a new type of reaction to be catalyzed by protein enzymes. Here we show how the alkenic amino acid vinylglycine can be generated in a biocompatible reaction from 5-allyloxy-2-amino-pent-3-enoate (APE) by ring-closing metathesis catalyzed by a standard Hoveyda-Grubbs catalyst. The vinylglycine produced in situ is then used as a precursor of isoleucine and methionine, thus allowing the growth of strains of Escherichia coli requiring these essential amino acids. The robust nutritional screen we have developed paves the way for the directed evolution of genetically encoded metathesis enzymes and the chemical elaboration of metathesis coenzymes.

## Introduction

Alkene metathesis stands out in the chemical industry as a versatile and straightforward reaction to reconnect C=C bonds, catalyzed by molybdenum VI or ruthenium II carbenoids (Schrock and Grubbs catalysts, respectively).[1–5] It seems to be completely absent from the catalytic arsenal and metabolism of extant living species. Therefore, its implementation *in vivo* would have the potential to augment biochemistry and to rewire a broad array of metabolic pathways. For the sake of illustration, the enzymatic conversion of two acrylate substrates into fumarate and ethylene might serve to redesign both central metabolism and hydrocarbon biosynthesis, while the reverse reaction (ethenolysis) might renew the bioproduction of acrylic monomers:

It is with such perspectives in mind that we set out to construct metabolic chassis and develop bacterial selection screens, in order to exploit metathesis enzymes developed by other research groups.[6] Acceleration of metathesis reactions by proteins in aqueous medium, albeit to modest rates, was pioneered by Ward’s and Hilvert’s laboratories.[7,8] These breakthroughs enabled the directed evolution of a “metathase” enzyme derived from streptavidin that uses a second generation Hoveyda-Grubbs ruthenium complex attached to a biotin handle for converting the substrate into a fluorescent umbelliferone product by ring-closing metathesis, with a turnover number higher than 600.[9]

Our goal was to set up a nutritional screen for selecting metathesis reactions that would be essential to bacterial cell. Such an approach amounts to implementing experimentally the paradigm of retrograde evolution of biosynthetic pathways[10] with the aim of mobilizing industrial catalysts *in vivo*. A few independent attempts to harness abiotic reactions for bacterial growth were described during the course of our study.[11,12] According to our plan, a bacterial strain would be genetically engineered such that one of its essential constituents would be produced through the conversion of a substrate provided exogenously in the medium together with a metathesis catalyst. The substrate conversion should occur in the aqueous medium surrounding bacterial cells and the product would then feed those cells that remain viable in presence of the metathesis reagent and catalyst. At a subsequent stage, such a vital metathesis reaction would serve to improve an enzyme like Ward’s “metathase”[9] secreted by bacteria and encoded from their genome by mutation and nutritional selection.

## Results and discussion

We targeted vinylglycine[13,14], the simplest amino acid bearing an alkene group, to become the product of a metabolic reaction of ring-closing metathesis. Vinylglycine was discovered long ago to occur naturally as its D-stereoisomer, which is produced by the fungus Rhodophyllus nidorosus.[15] Its L-isomer is known to be only mildly toxic to Escherichia coli and other living cells. The second-generation ruthenium catalyst HGII, which is commonly used in organic synthesis and remains active in presence of water[16], was chosen as “aproteic coenzyme” to obtain a proof of principle and show the generality of the approach.

Cystathionine synthase, the enzyme of methionine biosynthesis encoded by the *metB* gene, catalyzes the condensation of O-succinyl-homoserine (OSH) with cysteine to give cystathionine, which is next converted to homocysteine and finally to methionine.[17] Strains of *E. coli* unable to form OSH by deletion of the *metA* gene require methionine to grow. Vinylglycine was reported to replace OSH as an electrophilic substrate for the MetB enzyme and to sustain growth of these auxotrophic strains instead of methionine.[18] It is also known that auxotrophic strains of *E. coli*, which are unable to form the intermediate oxobutanoate from threonine and therefore require isoleucine as nutrient, can be grown using vinylglycine as nutrient instead of isoleucine or oxobutanoate. The enzyme tryptophan synthase, encoded by the *trpB* gene, is thought to catalyze the conversion of vinylglycine to oxobutanoate[19], whereas it is inactive toward threonine. The conversion of vinylglycine into the products cystathionine and oxobutanoate can both be rationalized according to the reaction mechanism of the pyridoxal phosphate coenzyme, pivoting on the quinonoid adduct.[18,19] We constructed several *E. coli* strains requiring an amino acid, namely a methionine auxotroph by deleting the gene *metA* from the chromosome (genotype *ΔmetA*) and an isoleucine auxotroph by deleting the *ilvA* gene and the tdc operon in the same strain (genotype *ΔilvA ΔtdcB*). These deletion markers were also combined to obtain a strain with a double nutritional requirement (*ΔilvA ΔtdcB ΔmetA*) auxotrophic for methionine and isoleucine (see Supporting Information, SI).

Growth assays were performed using radial gradients in plates of solid growth medium, a robust procedure used in microbiology to characterize nutrient or toxic agents such as antibiotics. Accordingly, a liquid suspension of bacterial cells (∼ 2 x 10^7^ cells in 5 mL) is spread and left to dry on the surface of the medium solidified with agar (volume 30 mL), then a small well (0.1 mL) is dug at the center of the plate and filled with a nutrient solution, which is allowed to diffuse into the agar gel during incubation (usually at 37°C). A positive response is visualized as an opaque disc of growing bacteria (Figure 1a). The growth area is proportional to the logarithm of the amount of nutrient introduced into the well.[20] Using this technique, we sought to demonstrate that a metathesis reaction could take place under biocompatible conditions and feed strains of *E. coli* by providing them with vinylglycine (Figure 1a). We first verified that the methionine auxotrophic strain *ΔmetA* displayed a vigorous growth ring on vinylglycine radial gradient plates after 24 h (Figure 2). The isoleucine auxotroph *ΔilvA ΔtdcB* showed a similar growth response to vinylglycine (Figure 2). These two positive results confirmed previous mention of the nutritional competence of vinylglycine[18] in the experimental format of radial gradients. The low toxicity of this compound is manifested in gradient plates by an inhibitory halo around the central well surrounded by a growth ring.

**Figure 1.**
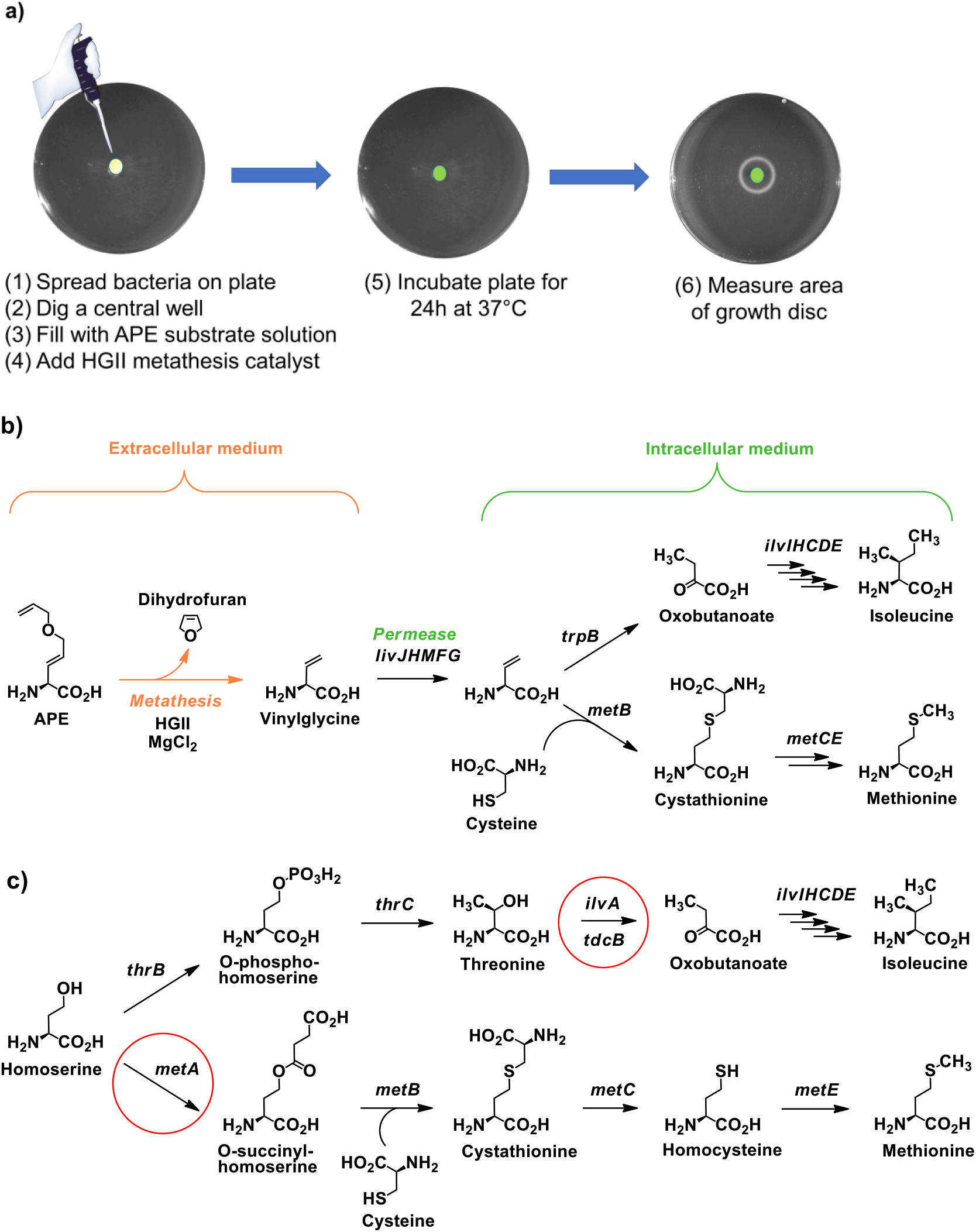
Biosynthesis of the essential amino acids isoleucine and methionine from vinylglycine generated by ring-closing metathesis in the extracellular medium. (a) Sketch of the experimental setup. Strains of *E. coli* strains auxotrophic for essential amino acids are incubated with the metathesis substrate APE together with the metathesis catalyst HGII/MgCl_2_ injected in a central well. Conversion of APE into vinylglycine by metathesis restores cell growth, as visualized by an opaque growth ring around the central well. (b) Composite pathways from APE to the end-products isoleucine and methionine, via vinylglycine, a known substrate of the enzymes tryptophan synthase (encoded by the *trpB* gene) and cystathionine synthase (encoded by *metB*). Intracellular metabolic steps are designated by the name of their corresponding genes. (c) Native pathways converting homoserine to isoleucine and methionine via O-phospho-homoserine and O-succinyl-homoserine, respectively. The metabolic steps inactivated by deletion of the *ilvA, tdcB* and *metA* genes in the isoleucine and methionine auxotrophic strains used to assay the conversion of APE by biocompatible metathesis are circled in red. APE, 5-allyloxy-2-amino-pent-3-enoate; *livJHMFG*, branched-chain amino acid ABC transporter; *trpB*, tryptophan synthase; *ilvIHCDE*, isoleucine-valine biosynthetic enzymes; *metB*, cystathionine synthase; *metC*, cystathionine beta-lyase; *metE*, homocysteine methyltransferase; *thrB*, homoserine kinase; *thrC*, threonine synthase; *ilvA*, threonine deaminase; *tdcB*, threonine deaminase; *metA*, homoserine succinyltransferase.

**Figure 2.**
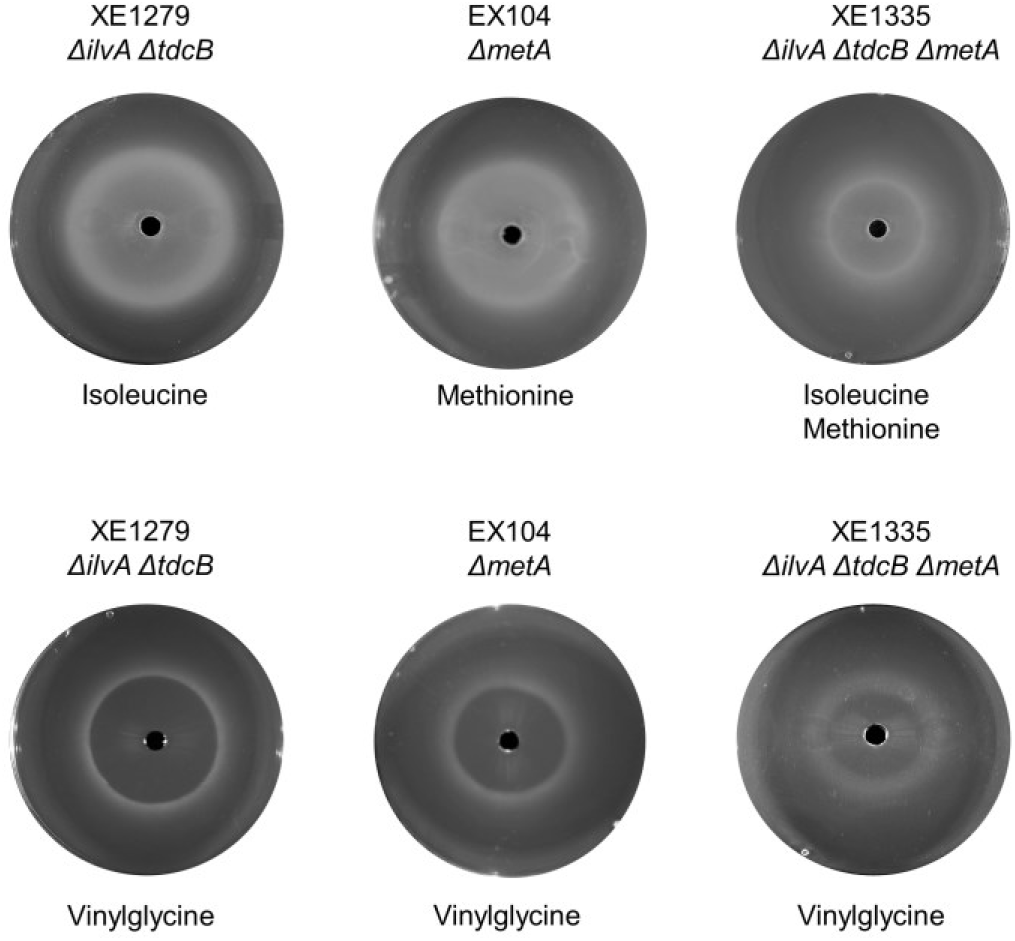
Nutritional assays of vinylglycine using radial gradient plates. (Top) Control plates supplied with 10 mM essential amino acids. (Bottom) Assay plates supplied with 10 mM vinylglycine.

The diene compound APE (L-stereoisomer of 5-allyloxy-2-amino-pent-3-enoic acid) was designed as a substrate for a ring-closing metathesis reaction to generate vinylglycine (Figure 1b). Benzyloxycarbonyl-protected L-vinylglycine was synthesized in six steps from L-methionine, as described by Rapoport et al. (SI).[21,22] Cross-metathesis with allyl alcohol was then performed[16] in fairly good yield (52% isolated) (SI). Further allylation of the alcohol moiety was attempted but failed, due to interfering reactivity of the α proton in the glycine moiety. Nevertheless, the desired compound could be obtained with a yield of 44% by palladium-catalyzed decarboxylative allylation (Figure 3) and finally deprotected into APE (SI).

**Figure 3.**
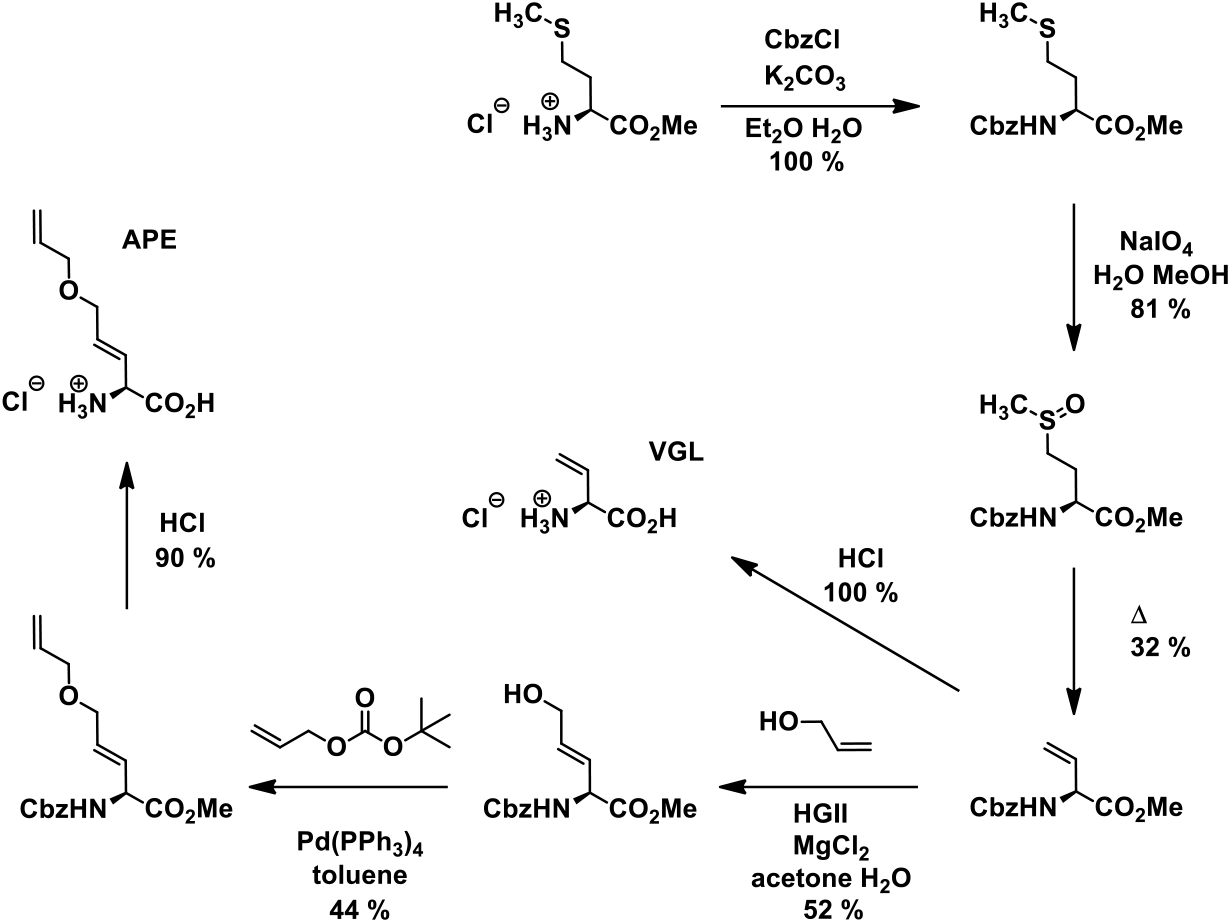
Synthesis of APE (L-stereoisomer of 5-allyloxy-2-amino-pent-3-enoic acid as its hydrochloride salt) by cross-metathesis between protected vinylglycine and allylic alcohol followed by palladium tetrakis-catalyzed O-allylation. The protocol for preparing L-vinylglycine hydrochloride (VGL), according to reference[21], is also shown. See Supplementary Information.

Strict care was taken to purify the synthetic intermediates in order to avoid the carry-over of trace amounts of nutrients. The APE compound thus prepared was not toxic to *E. coli* at any concentration. It did not support the growth of any auxotrophic strain, indicating that it was free from traces of methionine, its sulfoxides, vinylglycine and any other assimilable metabolites such as oxobutanoate or aminobutyrate.

Our main objective was to assess whether an off-the-shelf metathesis catalyst could be used to convert the APE substrate (Figure 1a) and augment the biosynthetic arsenal of *E. coli*. We therefore turned to the ruthenium complex HGII which is known to be active in acetone/water or tert-butanol/water mixtures.[23] Both solvents are known to be biocompatible, acetone being the least toxic to *E. coli*. Aqueous mixtures containing HGII, MgCl_2_ and either solvent were remarkably well tolerated by *E. coli*. Such aqueous solutions of the metathesis catalyst readily converted APE to vinylglycine at 37°C. The concentration of organic solvent affected critically catalysis by HGII, the production of vinylglycine decreasing steeply below 30/70 acetone/water (Figure 4).

**Figure 4.**
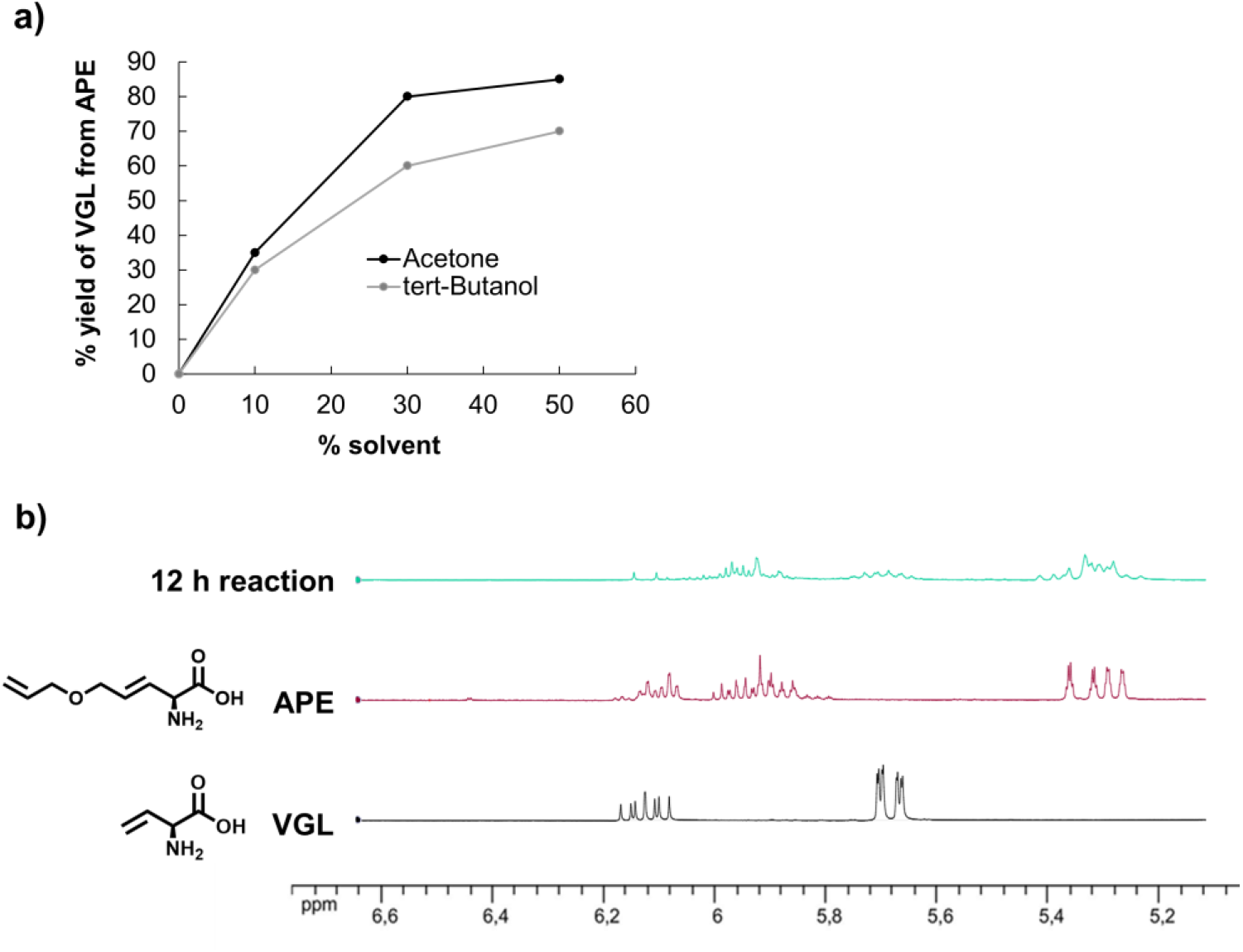
Optimisation of biocompatible metathesis conditions for converting APE to vinylglycine (VGL). (a) Yield of vinylglycine obtained in the presence of the metathesis catalyst in water/acetone and water/tert-butanol solutions at different concentrations (100/0, 90/10, 70/30 and 50/50). The 30/70 concentration of acetone/water is optimal for biocompatible vinylglycine production. (b) Course of the ring-closing metathesis reaction followed by 1H-NMR. See Supplementary Information (SI). The NMR spectra of APE and vinylglycine are shown in red and black, respectively; the spectrum of reaction intermediates is in green.

Having established (i) the nutritional potency of vinylglycine in radial gradients from a 30/70 acetone/water solution, (ii) effective metathesis reaction conditions in aqueous solvent mixtures, (iii) the innocuity of the APE metathesis substrate, (iv) the innocuity of the metathesis co-product dihydrofuran, (v) the innocuity of the common HGII catalyst towards *E. coli* cells, we then proceeded to assay biocompatible metathesis to generate vinylglycine and feed *E. coli* auxotrophs in radial gradient plates.

Subjecting the isoleucine auxotrophic strain *ΔilvA ΔtdcB* to a radial diffusion gradient of the reaction mix containing APE (1 eq), HGII (2 eq), MgCl_2_ (10 eq) in 30/70 acetone/water generated a growth disk corresponding to a calculated concentration of 1 to 3 mM vinylglycine (Figure 5). No bacterial growth was detectable when APE or HGII was omitted from the reaction mixture.

**Figure 5.**
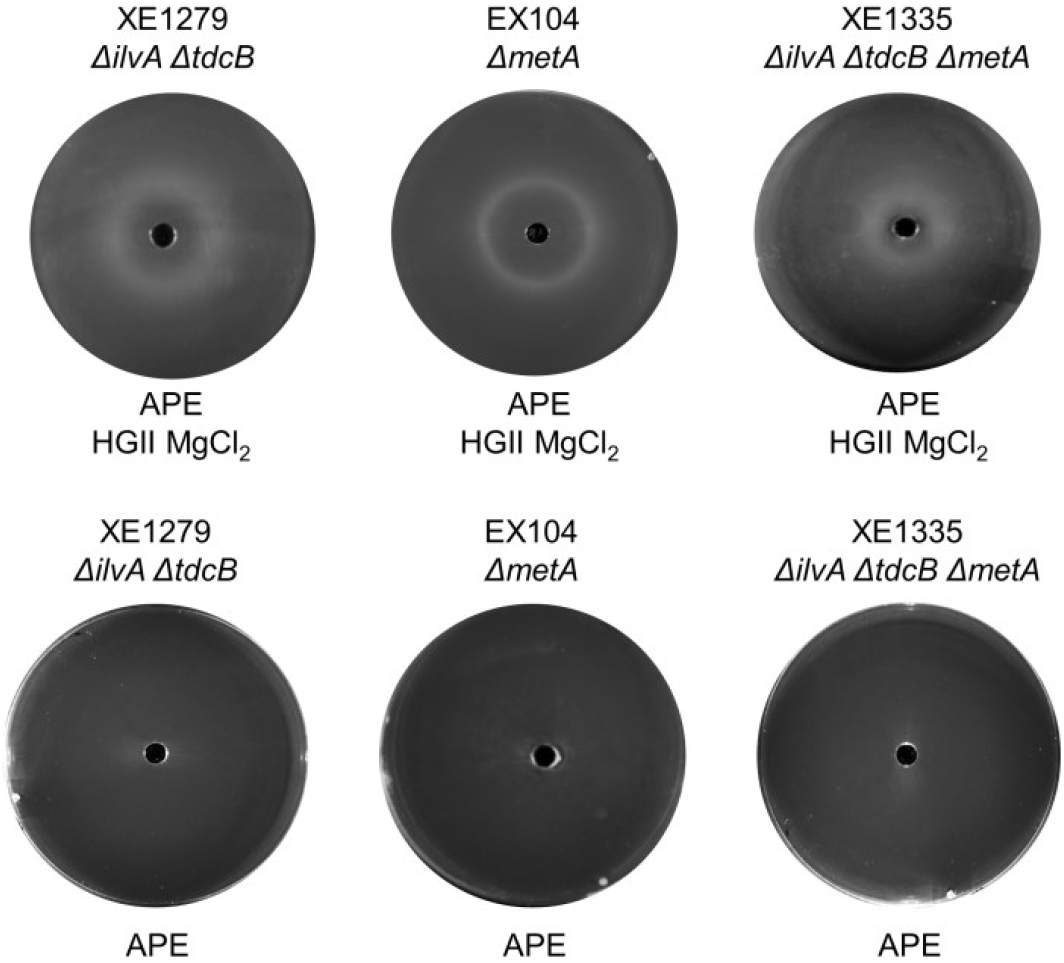
Nutritional utilization of APE as essential amino acid precursor through biocompatible metathesis. (Top) Radial gradients of the metathesis substrate APE together with the catalyst HGII/MgCl_2_ in 30/70 acetone/water solution allows growth of isoleucine and methionine auxotrophic strains, as visualized by an opaque ring around the central well. (Bottom) Control plates supplied with 70 mM APE and lacking the metathesis catalyst.

This result demonstrated that a metathesis reaction could support the production of an essential amino acid precursor and the concomitant growth of bacteria in the same medium. The methionine auxotroph *ΔmetA* gave a similar growth response to APE provided the HGII catalyst was present in the reaction mix at the center of the radial gradient. That vinylglycine generated by metathesis underlay the growth responses of isoleucine and methionine auxotrophs was further corroborated by the observation that the *ΔilvA ΔtdcB ΔmetA* strain requiring jointly the two amino acids could also grow on radial gradients of the reaction mix (Figure 5). In this case, the growth disc showed a reduction in size (28 mm diameter) compared to those observed for *ΔmetA* (37 mm) and *ΔilvA ΔtdcB* (47 mm), consistent with competition for vinylglycine between the biosynthetic pathways to isoleucine and methionine.

Overall, the rate of vinylglycine production in gradient assays seemed commensurate with enzyme kinetics sustaining biosynthesis in bacteria. The stoichiometry of the mixture was such that two equivalents of the HGII catalyst were present relative to the APE substrate. This substrate to catalyst ratio is small enough to justify the genetic selection of catalytic turnover at future stages of the investigation.

In conclusion, rapid advances in metabolic metathesis are to be expected from the chemical and microbiological constructs reported here. Our biocompatible reaction format should allow the structural exploration and functional comparison of metathesis catalysts[23–26] for metabolic purposes, with the priority of benchmarking the efficiency of ruthenium[16], molybdenum[2], tungsten[1], and even iron[27] in organometallic coenzymes to be developed next. Our tight nutritional screens should promote the evolutionary improvement of apoenzymes displayed at the surface of auxotrophic bacteria and combining with such coenzymes, supplied exogenously. Finally, the present study establishes vinylglycine as a non-canonical metabolite of great simplicity, reactivity and versatility for synthetic biology.

## Materials and methods

### Chemical methods

All reagents are commercial grade and were used as received. All moisture-sensitive reactions were carried out in oven-dried glassware under N_2_ Thin-layer chromatograms (TLC) and flash chromatography separations were respectively performed on precoated silica gel 60 F254 plates (Merck, 0.25 mm) and on silica gel 60 (230-400 mesh). ^1^H NMR spectra were recorded at 400 MHz. ^13^C NMR spectra were obtained at 100 MHz. High-resolution mass spectra HRMS were obtained from an LCT Premier XE using electrospray ionization coupled with a time flight analyser (ESI-TOF) and were performed by the Service de Spectometrie de Masse, IMAGIF/ICSN-CNRS, Gif-sur-Yvette, France or Institut de Chimie Organique et Analytique ICOA, Université d’Orléans, France. ESITOF (electrospray) mass spectra were measured on a Mariner Perspective Biosystem apparatus.

### Preparation of APE

#### N-[(benzyloxy)carbonyl]-L-methionine methyl ester

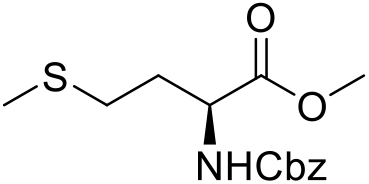

To a solution of L-methionine methyl ester hydrochloride (10.0 g, 50.0 mmol, 1 eq) and potassium carbonate (24.0 g, 173.0 mmol, 3.5 eq) in a biphasic system H_2_O/Et_2_O (50:50, 200 mL) at 0°C was added dropwise benzyl chloroformate (8.0 mL, 56.0 mmol, 1.1 eq). The mixture was warmed to R.T. and vigorously stirred for 12 h. The organic layer was separated, washed with hydrochloric acid (0.01 N), brine, dried over MgSO_4_, filtrated and concentrated to give N-[(benzyloxy)carbonyl]-L-methionine Methyl Ester (14.86 g, 100%).

#### NMR and HRMS data for the product

^**1**^**H NMR (CDCl**_**3**_): δ = 7.35-7.29 (m, 5H, Ar), 5.47 (d, J = 8.2 Hz, 1H, N**H**), 5.08 (s, 2H, COOC**H**_**2**_), 4.51-4.44 (m, 1H, CH_2_C**H**CO_2_Me), 3.72 (s, 3H, COOC**H**_**3**_), 2.50 (t, J = 7.3 Hz, 2H, SC**H**_**2**_CH_2_), 2.18-2.09 (m, 2H, SCH_2_C**H**_**2**_), 2.06 (s, 3H, C**H**_**3**_SCH_2_), 1,98-1.90 (m, 2H, SCH_2_C**H**_**2**_).

^**13**^**C NMR (CDCl**_**3**_): δ = 172.5 (**C**OOMe), 155.9 (N**C**OO), 136.2, 128.4, 128.1, 128.0, 66.9 (**C**H_2_OCO), 53.0 (**C**HCOOMe), 52.4 (COO**C**H_3_), 31.7 (CH**C**H_2_CH_2_), 29.8 (CHCH_2_**C**H_2_), 15.3 (S**C**H3).

**HRMS (ESI):** *m/z* calculated for C14H19NO4S, [M + H^+^] 298.1113, found 298.1109

#### Methyl L-2-[[(benzyloxy)carbonyl]amino]-4-(methylsulfinyl)butanoate

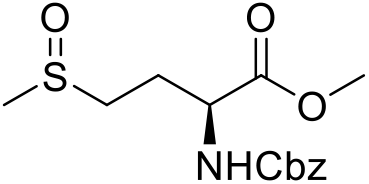

To an ice-cold solution of N-[(benzyloxy)carbonyl]-L-methionine methyl ester (14.82 g, 55.6 mmol, 1eq) in MeOH (120 mL) was added dropwise a solution of sodium metaperiodate (13.15 g, 61.0 mmol, 1.1 eq) in H_2_O (60 mL). The mixture was warmed to R.T. and stirred overnight and filtrated. The solid was washed with MeOH and the filtrate was concentrated to 80 mL. The aqueous layer was extracted with CH_2_Cl_2_ (3 X 100 mL). The organic layers were combined, washed with H_2_O, dried over MgSO_4_, filtrated and concentrated to give methyl L-2-[[(benzyloxy)carbonyl]amino]-4-(methylsulfinyl)butanoate (12.6 g, 81 %). The product was used without purification and presents different diastereoisomers.

##### NMR and HRMS data for the product

^**1**^**H NMR (CDCl**_**3**_): δ = 7.35-7.27 (m, 5H, Ar), 5.81 (dd, J_1_ = 25.3 Hz, J_2_ = 7.8 Hz, 1H, N**H**), 5.07 (s, 2H, COOC**H**_**2**_), 4.50-4.40 (m, 1H, CH_2_C**H**CO2Me), 3.73 (s, 3H, COOC**H**_**3**_), 2.80-2.62 (m, 2H, SC**H**_**2**_CH_2_), 2.50 (s, 3H, C**H**_**3**_SCH_2_), 2.40-2.28 (m, 1H, SCH_2_C**H**_**2**_), 2.19-2.04 (m, 1H, SCH_2_C**H**_**2**_).

^**13**^**C NMR (CDCl**_**3**_): δ = 171.9 (**C**OOMe), 156.2 (N**C**OO), 136.0, 128.7, 128.4, 128.3, 67.3 (**C**H_2_OCO), 53.2 (**C**HCOOMe), 52.9 (COO**C**H_3_), 50.4 (CHCH_2_**C**H_2,_ D1), 50.3 (CHCH_2_**C**H_2,_ D2), 38.7 (S**C**H3), 26.2 (CH**C**H_2_CH_2,_ D1), 25.9 (CH**C**H_2_CH_2,_ D2).

**HRMS (ESI):** *m/z* calculated for C14H19NO5S, [M + H^+^] 314.1062, found 315.1068

#### N-[(benzyloxy)carbonyl]-L-vinylglycine methyl ester

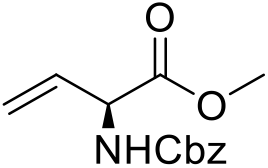

Methyl L-2-[[(benzyloxy)carbonyl]amino]-4-(methylsulfinyl)butanoate (12.6 g, 40.0 mmol, 1 eq) was distillated at 190 °C (15 mBar). The crude oil was purified by flash chromatography on silica gel (Heptane/AcOEt, 70:30) to give N-[(benzyloxy)carbonyl]-L-vinylglycine methyl ester (3.2 g, 32 %).

##### NMR and HRMS data for the product

^**1**^**H NMR (CDCl**_**3**_): δ = 7.37-7.27 (m, 5H, Ar), 5.93-5.84 (m,1H, CH_2_=C**H**), 5.63 (d, J = 7.5 Hz, 1H, N**H**), 5.34 (dd, J_1_ =17.2 Hz, J_2_ = 1.5 Hz, 1H, C**H**_2_=CH), 5.25 (dd, J_1_ =10.3 Hz, J_2_ = 1.5 Hz, 1H, C**H**_2_=CH), 5.10 (s, 2H, COOC**H**_**2**_), 4.94-4.91 (m, 1H, CHC**H**CO_2_Me), 3.73 (s, 3H, COOC**H**_**3**_).

^**13**^**C NMR (CDCl**_**3**_): δ = 171.0 (**C**OOMe), 155.7 (N**C**OO), 136.3, 132.4 (CH_2_=**C**H), 128.6, 128.3, 128.2, 127.0, 117.9 (**C**H_2_=CH), 67.2 (**C**H_2_OCO), 56.2 (**C**HCOOMe), 52.8 (COO**C**H_3_).

**HRMS (ESI):** *m/z* calculated for C13H15NO4, [M + Na^+^] 272.0899, found 272.0891.

#### L-vinylglycine hydrochloride salt

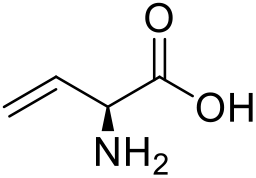

A solution of N-[(benzyloxy)carbonyl]-L-vinylglycine methyl ester (1.00 g, 4.0 mmol, 1 eq) in HCl 6N (3 mL) was refluxed for 4 h and concentrated. The residue was washed with Et_2_O to give L-vinylglycine hydrochloride (VGL) (0.546 g, 100%).

##### NMR and HRMS data for the product

^**1**^**H NMR (D**_**2**_**O):** δ = 6.04-5.95 (m, 1H, CH_2_=C**H**), 5.56 (dd, J_1_= 3.4 Hz, J_2_= 1.2 Hz, 1H, C**H**_**2**_=CH), 5.53 (dd, J_1_= 3.2 Hz, J_1_= 1.2 Hz, 1H, C**H**_**2**_=CH), 4.49 (d, J =7.3 Hz, 1H, CH_2_=CHC**H**NH_2_).

^**13**^**C NMR (D**_**2**_**O):** δ = 170.3 (**C**OOH), 127.7 (CH_2_=**C**H), 123.1 (**C**H_2_=CH), 54.9 (**C**HCOOMe).

**HRMS (ESI):** *m/z* calculated for C4H7NO2, [M + H^+^] 102.0555, found 102.0552.

#### (S,E)-methyl 2-(((benzyloxy)carbonyl)amino)-5-hydroxypent-3-enoate

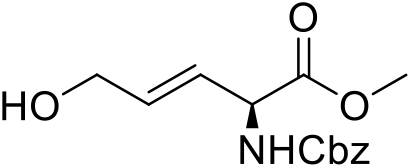

To a solution of N-[(benzyloxy)carbonyl]-L-vinylglycine methyl ester (1.00 g, 4.0 mmol, 1 eq) and MgCl_2_.6H_2_O (0.81 g, 4.0 mmol, 1 eq) in *t*-BuOH (30 mL) and H_2_O (30 mL) was added Hoveyda Grubbs II (0.25 g, 0.2 mmol, 0.05 eq). Allyl alcohol (2.32 g, 40.0 mmol, 10 eq) was added dropwise. The mixture was stirred overnight and concentrated under vacuum. The residue was purified by silica gel chromatography (Heptane/Ethyl acetate (60:40 to 30:70)) to give (S,E)-methyl 2-(((benzyloxy)carbonyl)amino)-5-hydroxypent-3-enoate as a yellow oil (0.585 g, 52%).

##### NMR and HRMS data for the product

^**1**^**H NMR (CDCl**_**3**_): δ = 7.35-7.24 (m, 5H, Ar), 5.93 (dt, J_1_ = 15.9 Hz, J_2_ = 4.5 Hz, 1H, CH_2_C**H**=CH), 5.77 (dd, J_1_ =15.9 Hz, J_2_ = 5.9 Hz, 1H, CH_2_CH=C**H**), 5.49 (d, J = 7.0 Hz, N**H**), 5.10 (s, 2H, COOC**H**_**2**_), 4.94-4.91 (m, 1H, CHC**H**CO_2_Me), 4.16 (d, J = 4.5 Hz, 2H, OH**C**H_2_), 3.75 (s, 3H, COOC**H**_**3**_).

^**13**^**C NMR (CDCl**_**3**_): δ = 171.2 (**C**OOMe), 155.8 (N**C**OO), 136.2, 133.2 (CH_2_CH=**C**H), 128.7, 128.4, 128.3, 125.0 (CH_2_**C**H=CH), 67.3 (**C**H_2_OH), 62.5 (**C**H_2_OCO), 55.5 (**C**HCOOMe), 53.0 (COO**C**H_3_).

**HRMS (ESI):** *m/z* calculated for C14H17NO5, [M + Na^+^] 302.0926, found 302.1002.

#### (S,E)-methyl 5-(allyloxy)-2-(((benzyloxy)carbonyl)amino)pent-3-enoate

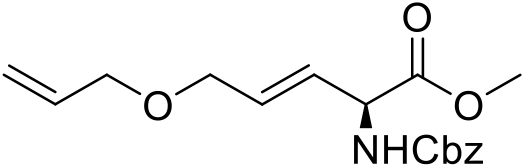

Allyl tertbutyl carbonate (0.29 g, 1.81 mmol, 1.6 eq), (S,E)-methyl 2-(((benzyloxy)carbonyl)amino)-5-hydroxypent-3-enoate (0.32 g, 1.14 mmol, 1 eq) and Pd(PPh_3_)_4_ (0.071 g, 0.06 mmol, 0.05 eq) were mixed in toluene (6 mL) and stirred for 4 hours, filtrated on celite and concentrated. The residue was purified by silica gel chromatography (Heptane/Ethyl acetate (70:30) to give (S,E)-methyl5-(allyloxy)-2-(((benzyloxy)carbonyl)amino)pent-3-enoate as a colorless oil (0.162 g, 44%).

##### NMR and HRMS data for the product

^**1**^**H NMR (CDCl**_**3**_): δ = 7.37-7.28 (m, 5H, Ar), 5.93-5.75 (m, 3H, CH_2_=C**H**, OCH_2_CH=C**H**, OCH_2_C**H**=CH), 5.44 (d, J = 7.6 Hz, N**H**), 5.25 (dd, J_1_ =17.4 Hz, J_2_ = 1.6 Hz, 1H, C**H**_2_=CH), 5.17 (dd, J_1_ =10.7 Hz, J_2_ = 1.6 Hz, 1H, C**H**_2_=CH), 5.10 (s, 2H, COOC**H**_**2**_), 4.94-4.91 (m, 1H, CHC**H**CO_2_Me), 3.98-3,94 (m, 4H, O**C**H_2_), 3.74 (s, 3H, COOC**H**_**3**_).

^**13**^**C NMR (CDCl**_**3**_): δ = 171.1 (**C**OOMe), 155.7 (N**C**OO), 136.3, 134.6 (CH_2_=**C**HCH), 130.5 (CH_2_=**C**HCH_2_), 128.7, 128.4, 128.3, 126.5 (CH_2_**C**H=CH), 117.2 (**C**H_2_=CH), 71.6 (O**C**H_2_CH=CHCH), 69.5 (CH_2_=CH**C**H_2_O), 67.3 (**C**H_2_OCO), 55.5 (**C**HCOOMe), 52.9 (COO**C**H_3_).

**HRMS (ESI):** *m/z* calculated for C17H21NO5, [M + Na^+^] 342.1317, found 342.1310.

#### (S,E)-5-(allyloxy)-2-aminopent-3-enoic acid hydrochloride salt

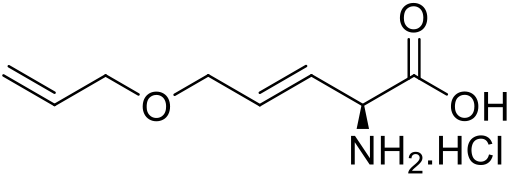

A solution of methyl 5-(allyloxy)-2-(((benzyloxy)carbonyl)amino)pent-3-enoate (0.81 g, 0.25 mmol, 1eq) in HCl 6N (1mL) was refluxed for 2h and concentrated. The residue was washed with Et_2_O to give (S,E)-5-(allyloxy)-2-aminopent-3-enoic acid hydrochloride salt (0.051 g, 90 %).

##### NMR and HRMS data for the product

^**1**^**H NMR (D**_**2**_**O):** δ = 6.15-6.06 (m, 1H, CH_2_=C**H**CH_2_), 6.01-5.85 (m, 2H, OCH_2_CH=C**H** + OCH_2_C**H**=CH), 5.34 (ddq, J_1_ =17.5 Hz, J_2_ = 1.6 Hz, 1H, C**H**_2_=CH), 5.28 (dq, J_1_ =10.7 Hz, J_2_ = J_3 =_ 1.6 Hz, 1H, C**H**_2_=CH), 4.51 (d, J =8.8 Hz, 1H, CH=CHC**H**NH_2_), 4.13 (dt, J_1_ =5.5 Hz, J_2_ = 0.6 Hz, 2H, O**C**H_2_CH=CH), 4.07 (dt, J_1_ =5.9 Hz, J_2_ = 1.6 Hz, 2H, CH_2_CHC**H**_**2**_OCH_2_), 3.76 (s, 3H, COOC**H**_**3**_).

^**13**^**C NMR (D**_**2**_**O):** δ = 170.7 (**C**OOH), 134.8 (CH=**C**HCH), 133.4 (CH_2_=**C**HCH_2_), 122.8 (**C**H=CHCH), 118.6 (**C**H_2_=CHCH_2_), 71.2 (O**C**H_2_CHCH), 68.8 (CH_2_CH**C**H_2_O), 54.4 (**C**HCOOMe).

**HRMS (ESI):** *m/z* calculated for C8H13NO3, [M + H^+^] 172.0974, found 172.0967.

### Ring-closing metathesis assays

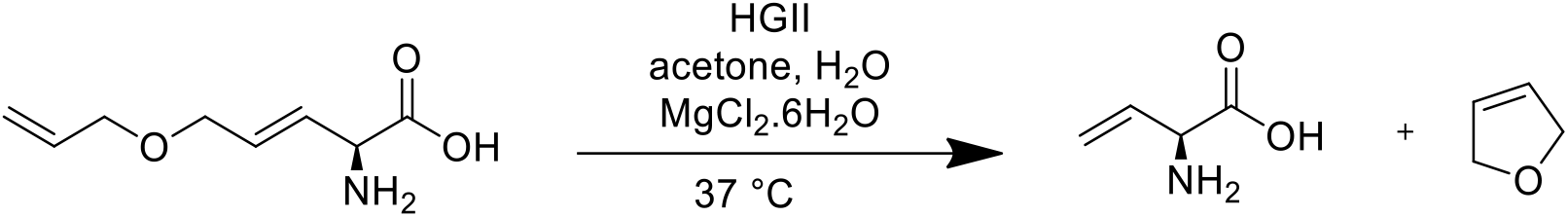

To a solution of N-[((S,E)-5-(allyloxy)-2-aminopent-3-enoic acid hydrochloride salt (0.014 g, 0.08 mmol, 1eq) and MgCl_2_.6H_2_O (0.018 g, 0.08 mmol, 1 eq) in acetone (0.3 mL) and H_2_O (0.7 mL) was added Hoveyda Grubbs II (0.011 g, 0.02 mmol, 0.2 eq). The mixture was stirred at 37 °C and followed by LCMS. Vinylglycine peak appears after 1h of reaction and the conversion is about 80% after 12 h. Vinylglycine peaks can also be observed by NMR after 4h of reaction.

### Biological methods

#### Growth media

Bacteria were grown in LB medium (Conda) or in minimal standard medium (MS)[28] supplemented with D-glucose (2 g/L). Growth media were solidified with 15 g/L agar for the preparation of Petri plates. Liquid and solid cultures were incubated at 37°C. When required, antibiotics were added at the following concentrations: carbenicillin, 100 mg/L (Invitrogen); spectinomycin (Sigma), 100 mg/L; apramycin (Sigma), 50 mg/L. For methionine or isoleucine auxotrophic strains, L-methionine (Sigma) or L-isoleucine (Sigma) was added to the culture medium at 0.1 mM concentration.

#### Construction of *E. coli* strains

The strains used and constructed in this study are all derivatives of the wild type *Escherichia coli* K12 strain MG1655 and are reported in table S1. To delete genes from the *E. coli* chromosome, a DNA fragment encoding a resistance gene (*aad* for Spectinomycin resistance, *aac* for Apramycine, *kan* for Kanamycin or *cat* for Chloramphenicol resistance) surrounded by 50 bp homologous to the chromosomal regions upstream and downstream of the gene or gene operon to be deleted was generated by PCR.

PCR reactions were carried out in 96-well microplates in 50 μl reactions containing 2.5 U of *TaKaRa Ex Taq* polymerase, 1 pg of the resistance cassette gene, 1.0 μM of each primer (X1004 and X1005 for *tdc* operon deletion, X1050 and X1051 for *ilvA*, and X1054 and X1055 for *metA*) and 200 μM dNTPs. Reactions were run for 30 cycles: 95°C for 30 s, 55°C for 30 s, 72°C for 2 min, plus an additional 2 min at 72°C.

The gene or operon to delete was then replaced by the antibiotic resistance cassette by homologous recombination using the lambda red recombinase system.[29] Strains, carrying the pKD46 plasmid coding for the lambda red recombinase, were grown in 10 ml LB cultures with ampicillin and L-arabinose at 30°C to an OD600 of ≈0.9 and then made electrocompetent by concentrating 100-fold and washing three times with cold H_2_O. 50 μl of competent cells were mixed with 500 ng of the PCR fragment in an ice-cold 0.2 cm cuvette (Bio-Rad Inc.). Cells were electroporated at 2.5 kV with 25 mF and 200 Ω, immediately followed by the addition of 1 ml of LB medium (2% Bacto Tryptone (Difco), 0.5% yeast extract (Difco), 10 mM NaCl, 2.5 mM KCl, 10 mM MgCl_2_, 10 mM MgSO_4_, 20 mM glucose). After incubation for 3h at 37°C, bacteria were spread onto agar plate to select CmR, ApraR or KmR recombinants.

To remove the antibiotic resistance cassettes, bacteria were transformed with pCP20 plasmid encoding the FLP recombinase.[29] Eight transformants were plated on LB and incubated overnight at 30°C to allow FLP recombinase expression. Cells were then re-streaked on LB at 42°C to eliminate the plasmid and screened for antibiotic sensitivity. The removal of the antibiotic resistance cassette was confirmed by PCR using the external oligonucleotides X1052 and X1053 for the *ΔilvA* locus, X1056 and X0157 for the *ΔmetA* locus, and X1006 and X1007 for the *ΔtdcABCDEFG* locus. All oligonucleotides used in this work are reported in Table S2.

**Table S1.** Bacterial strains used in the study.

| Name | Genotype |
| --- | --- |
| MG1655 | *E. coli* K12 (wt) |
| EX104 | *E. coli* K12 $\Delta metA::cat$ |
| XE 1279 | *E. coli* K12 $\Delta tdcABCDEFG::aad$ $\Delta ilvA::aac$ |
| XE 1335 | *E. coli* K12 $\Delta tdcABCDEFG$ $\Delta ilvA$ $\Delta metA::kan$ |

**Table S2.**
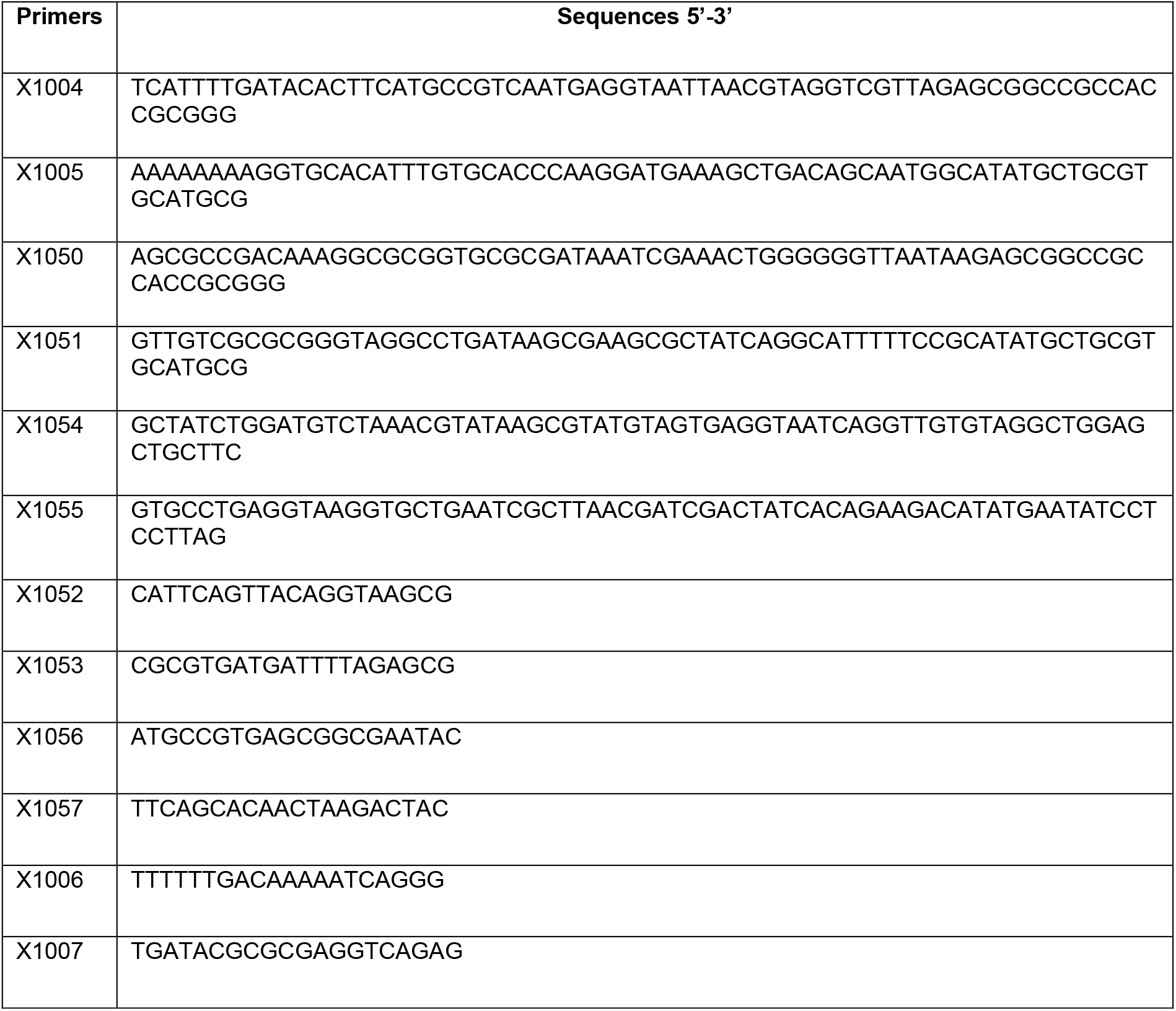
Oligonucleotides used in this work.

#### Nutrition assays using radial gradients

Bacteria were grown overnight in MS glucose liquid cultures, centrifuged (for 3 min at 7,000 rpm) and rinsed 3 times with fresh MS medium, diluted to an OD (600 nm) of 0.04 in 5 mL of fresh medium and spread on plates of the same medium solidified with agar. A volume of 4.5 mL excess liquid was removed by suction, the surface of the plates briefly left to dry, a central well was dug and 80 μL of an aqueous solution of isoleucine, methionine or vinylglycine at different concentrations were injected in the central well. Plates were incubated overnight at 37°C. After 24 h, the plates were photographed under intense white light. Absence of growing bacteria appeared as a grey background and growing bacteria as a white ring.

#### Toxicity assays using radial gradients

Bacteria were grown overnight in MS glucose liquid cultures containing methionine or isoleucine at 300 μM concentration, centrifuged (for 3 min at 7,000 rpm) and rinsed 3 times with fresh MS medium, diluted to an OD of 0.04 in 5 mL of fresh medium and spread on plates of the same medium solidified with agar. A volume of 4.5 mL excess liquid was removed by suction and the surface of the plates briefly left to dry. A central well was dug in the agar and 80 μL of an aqueous solution of APE, HGII, vinylglycine or dihydrofuran was injected in the well. The plates were incubated at 37°C for 24 h. Then, photographs were taken under intense white light. Absence of growing bacteria appeared as a halo of grey background and growing bacteria as a white corona.

#### Metabolic metathesis assays using radial gradients

Bacteria were grown overnight in MS glucose liquid cultures containing methionine or isoleucine at 100 μM concentration, centrifuged (for 3 min at 7,000 rpm) and rinsed 3 times with fresh MS medium, diluted to an OD of 0.04 in 5 mL of fresh medium and spread on plates of the same medium solidified with agar. A volume of 4.5 mL excess liquid was removed by suction and the surface of the plates briefly left to dry. A central well was dug at the center of the plates. Then, 56 μL of an aqueous solution of (E)-5-(allyloxy)-2-aminopent-3-enoic acid (APE, 100 mM) and MgCl_2_.6H_2_O (0.014g, 10eq) were injected in the well, followed by 24 μL of a saturated solution of HGII (0.008 g, 2 eq) in acetone. The plates were incubated at 37°C for 24 h. Then, photographs were taken under intense white light.

## Supporting information

NMR spectra

---

**Figure S1.**
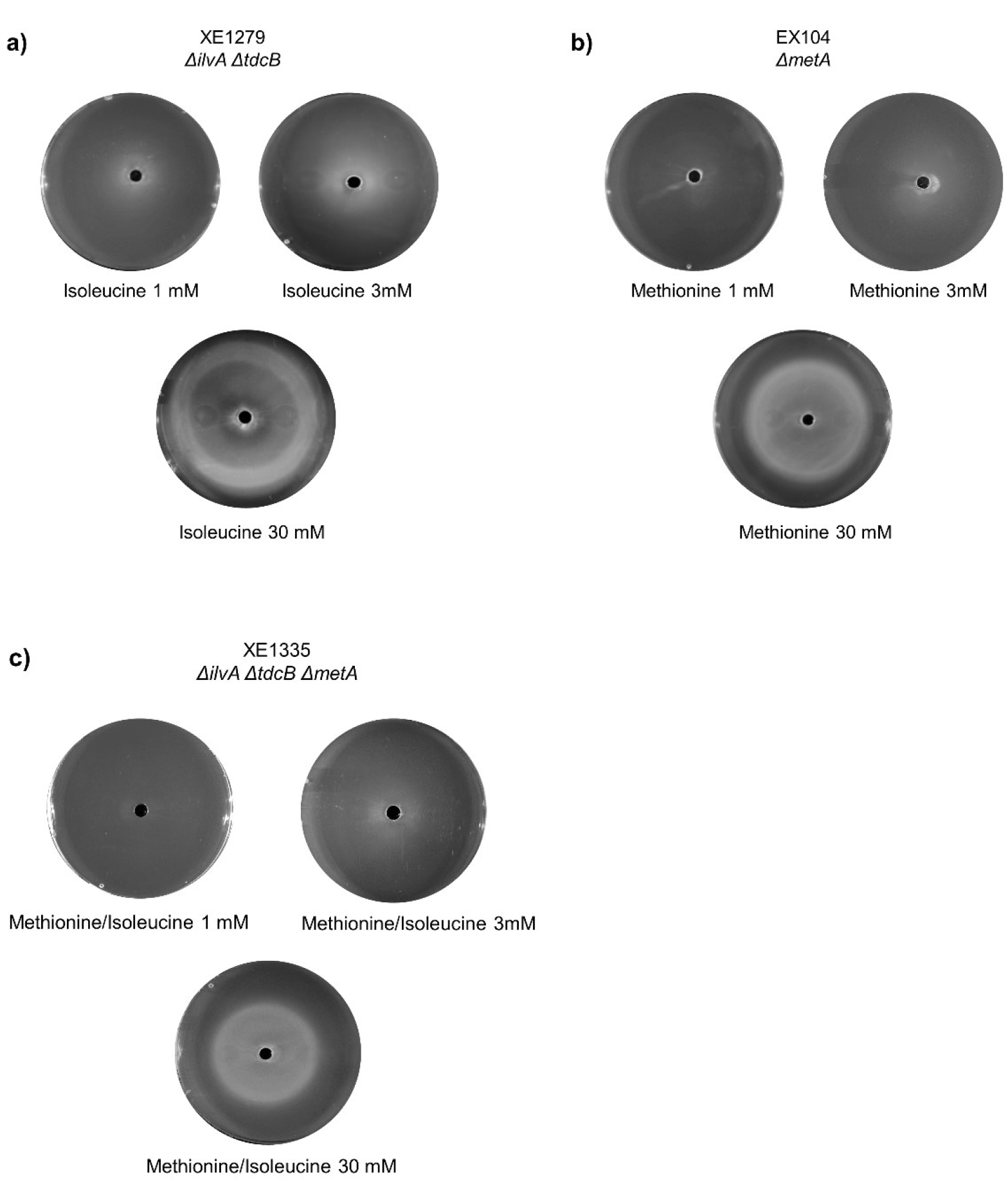
Calibration. Growth patterns of strains auxotrophic for isoleucine (a), methionine (b), and both amino acids (c) on radial gradient plates containing different amino acids diffusing from a central well.

**Figure S2.**
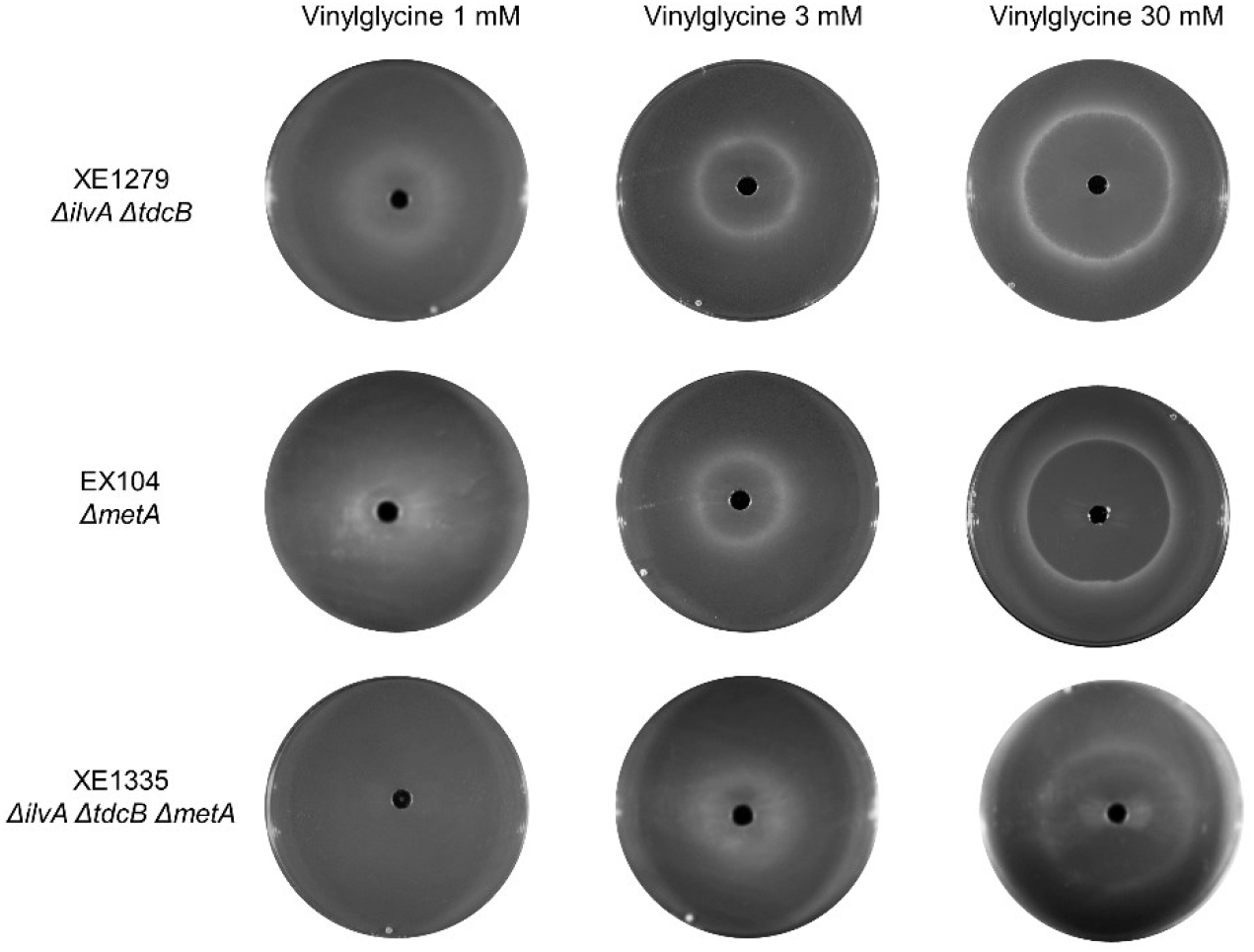
Growth patterns of strains auxotrophic for isoleucine (XE1279), methionine (EX104), and both amino acids (XE1335) on radial gradient plates of vinylglycine diffusing from a central well.

**Figure S3.**
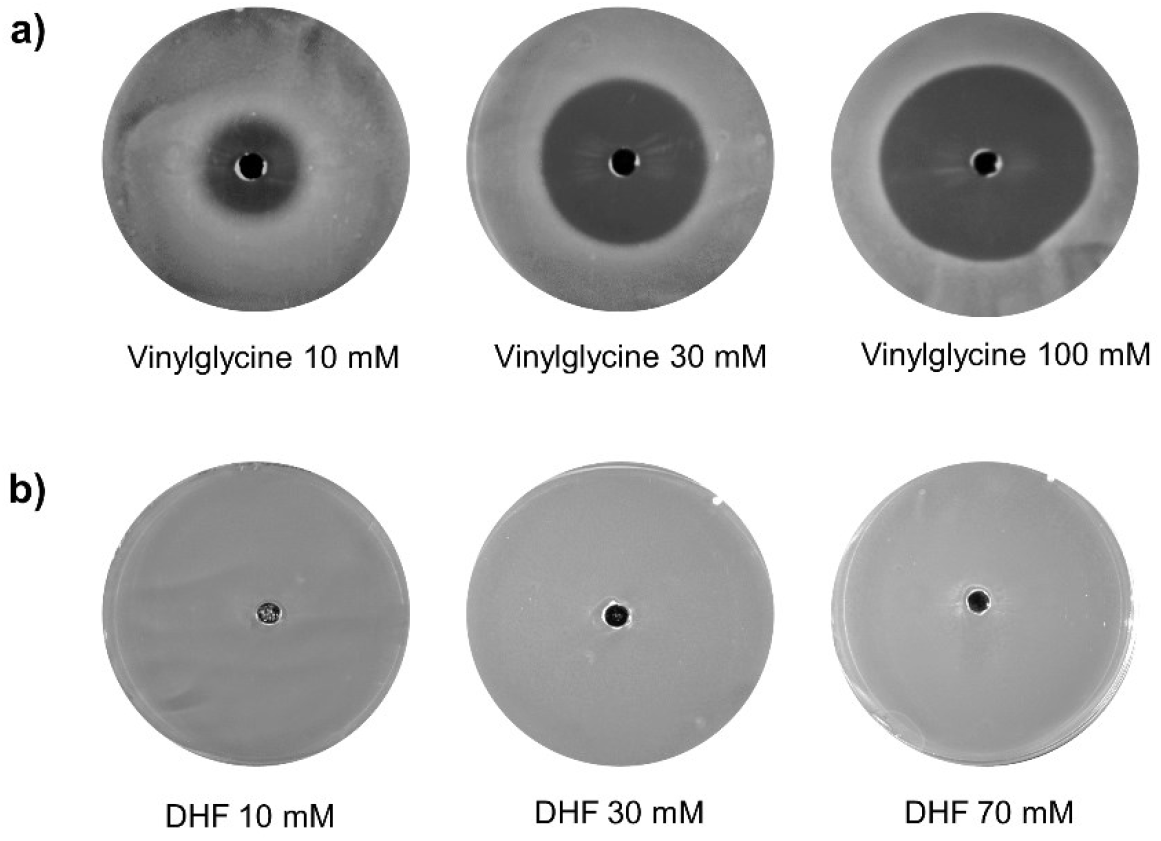
Toxicity toward the isoleucine auxotrophic strain XE1279. **a)** Vinylglycine showed a slight toxicity. **b)** Dihydroxyfuran (DHF), the coproduct of the metathesis reaction, showed no toxicity.

**Figure S4.**
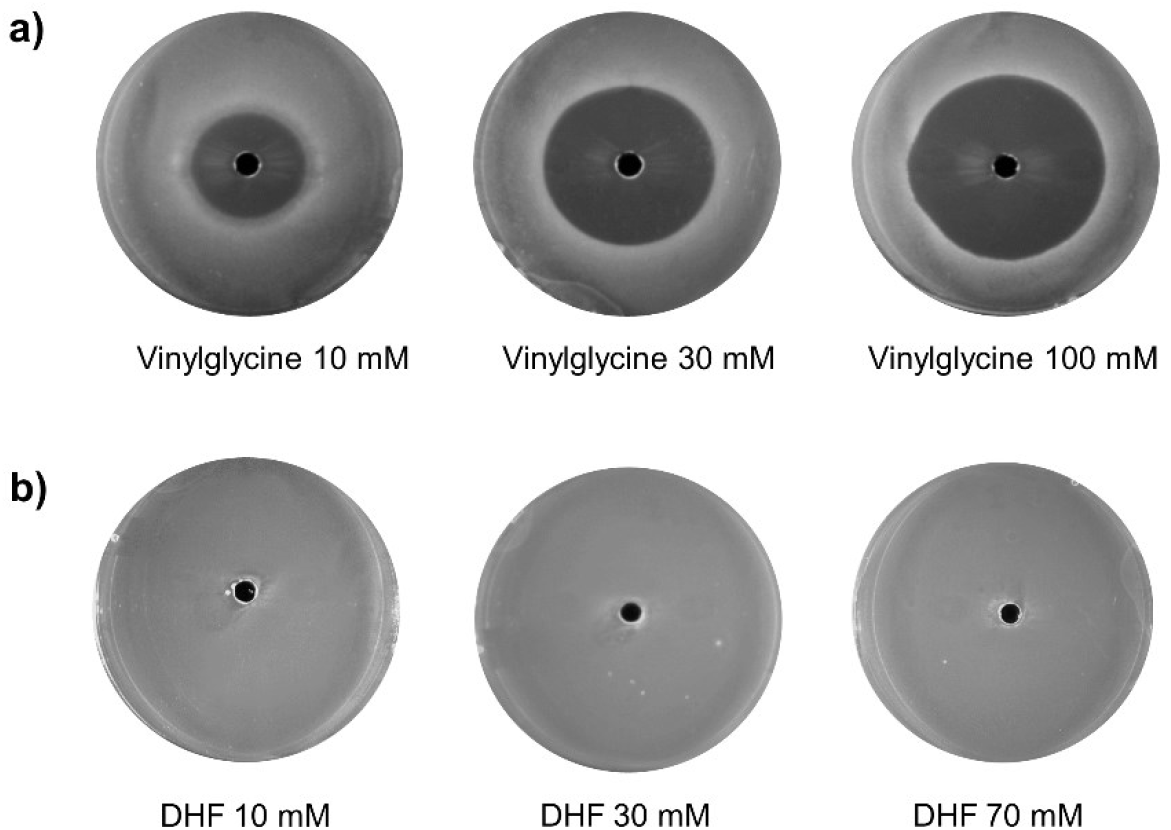
Toxicity toward the methionine auxotrophic strain EX104. **a)** Vinylglycine showed a slight toxicity. **b)** Dihydroxyfuran (DHF), the coproduct of the metathesis reaction, showed no toxicity.

**Figure S5.**
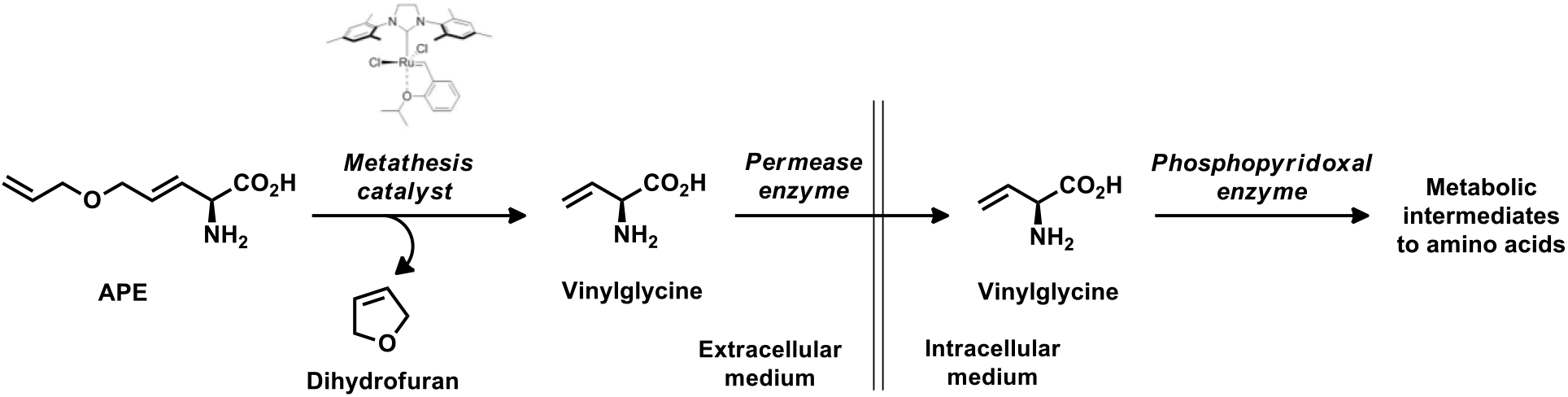
Metabolic metathesis scheme

## Acknowledgements

This work was funded by CEA, Genopole, Université d’Evry, CNRS and TESSSI. We are grateful to Dr. Alessandro De Simone for his help in writing and documenting the manuscript. We also thank Dr. Jean-Marc Paris, Prof. Janine Cossy and Prof. Jean-Marie Lehn for discussions.

## Notes

### Competing Interest Statement

The authors have declared no competing interest.

