## Supplementary figures and images for "An emergent biosynthetic pathway to essential amino acids by metabolic metathesis"

### NMR spectra

**NMR spectra**

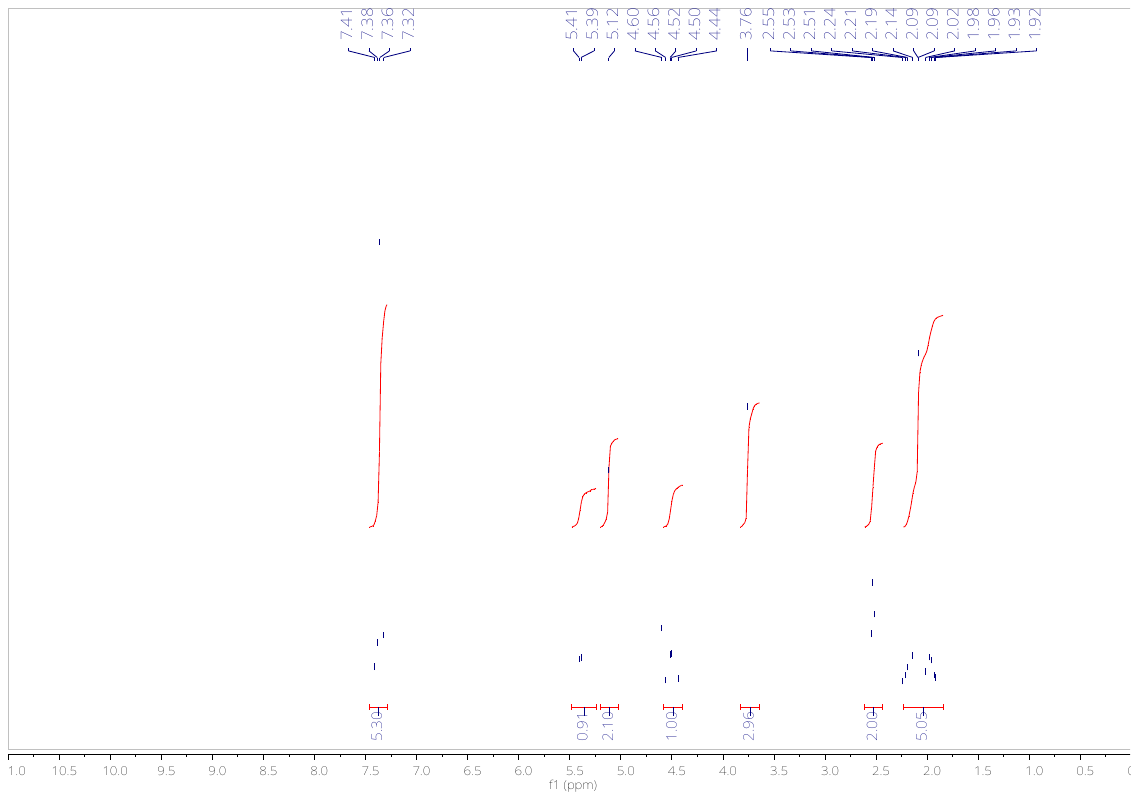

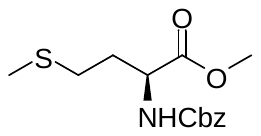

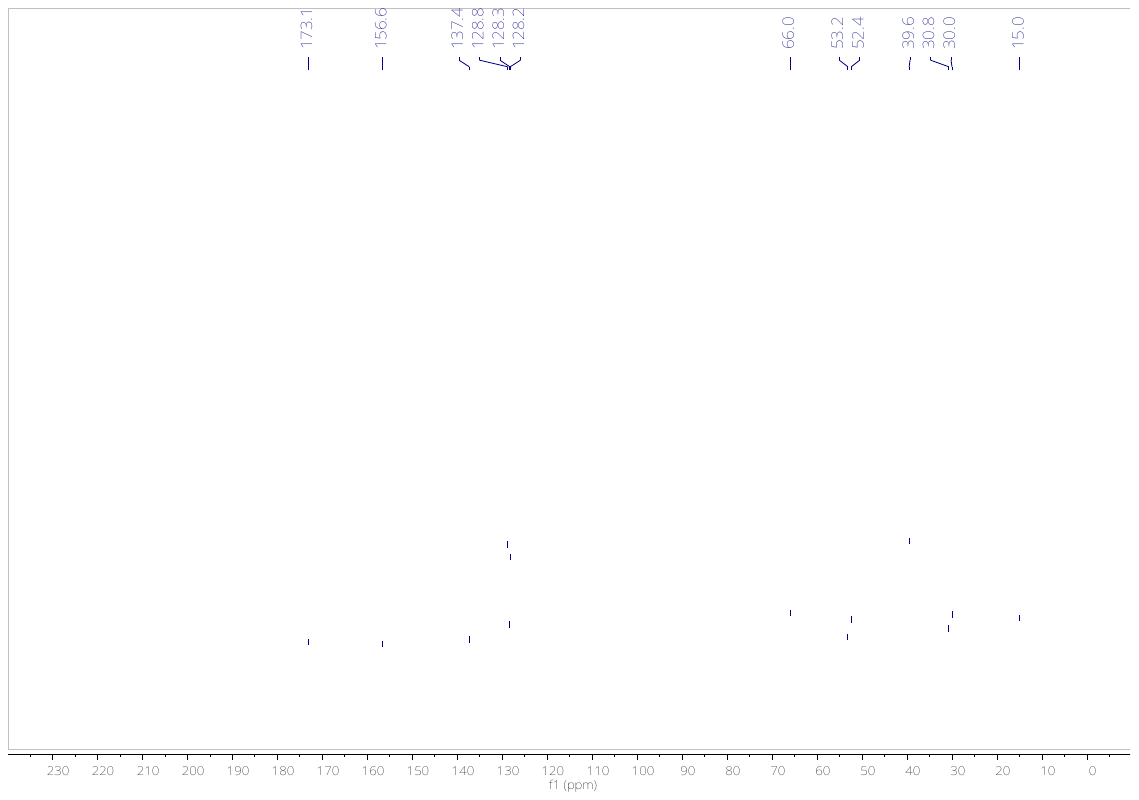

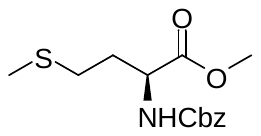

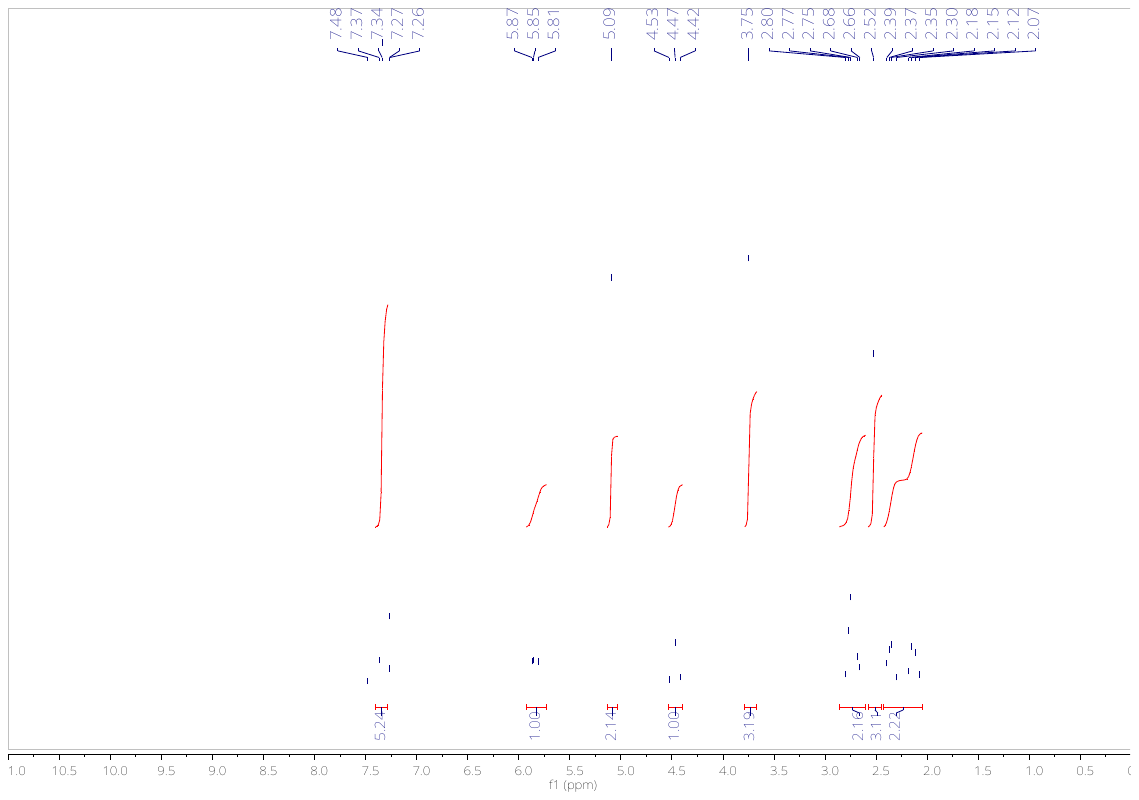

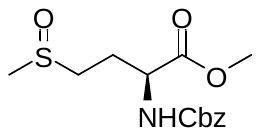

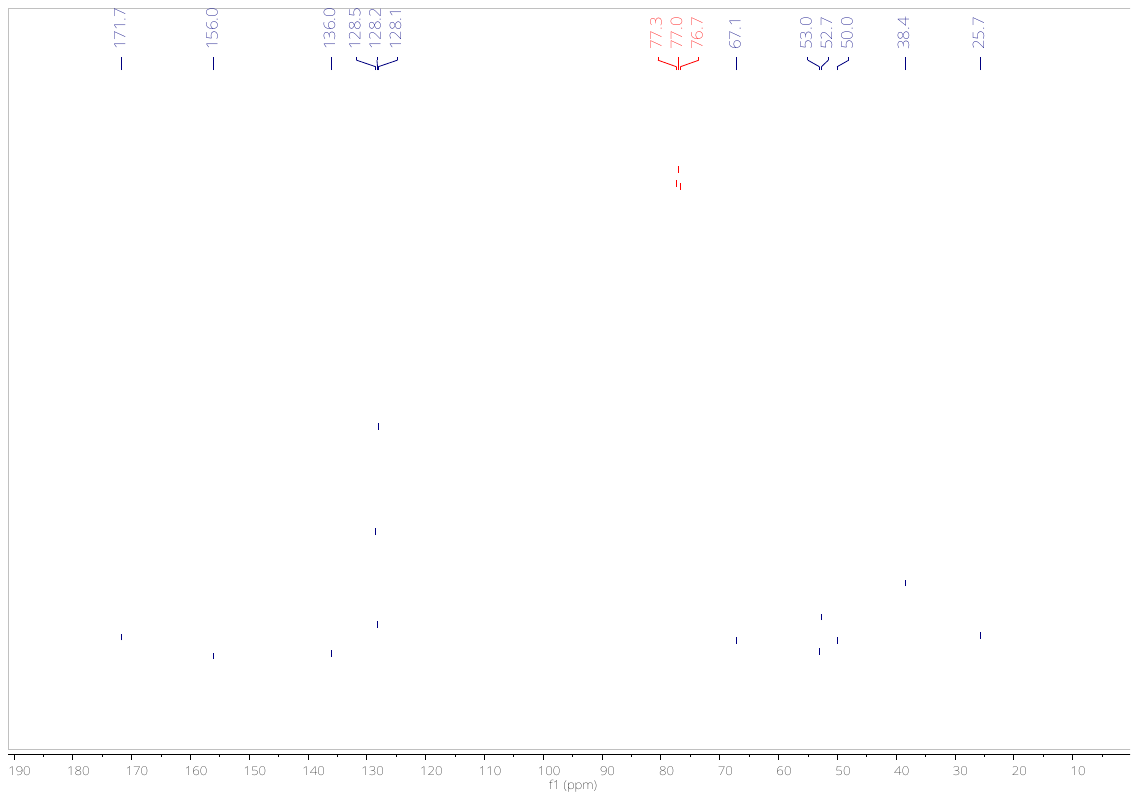

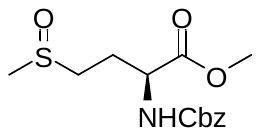

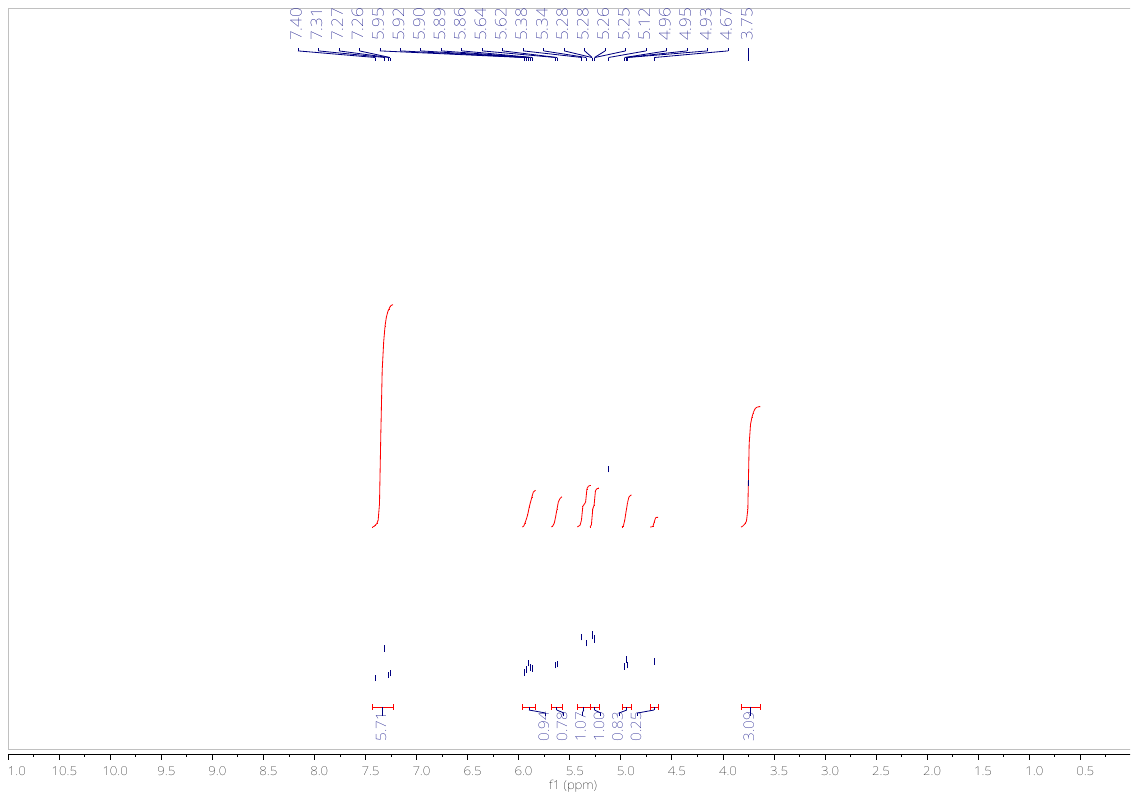

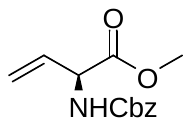

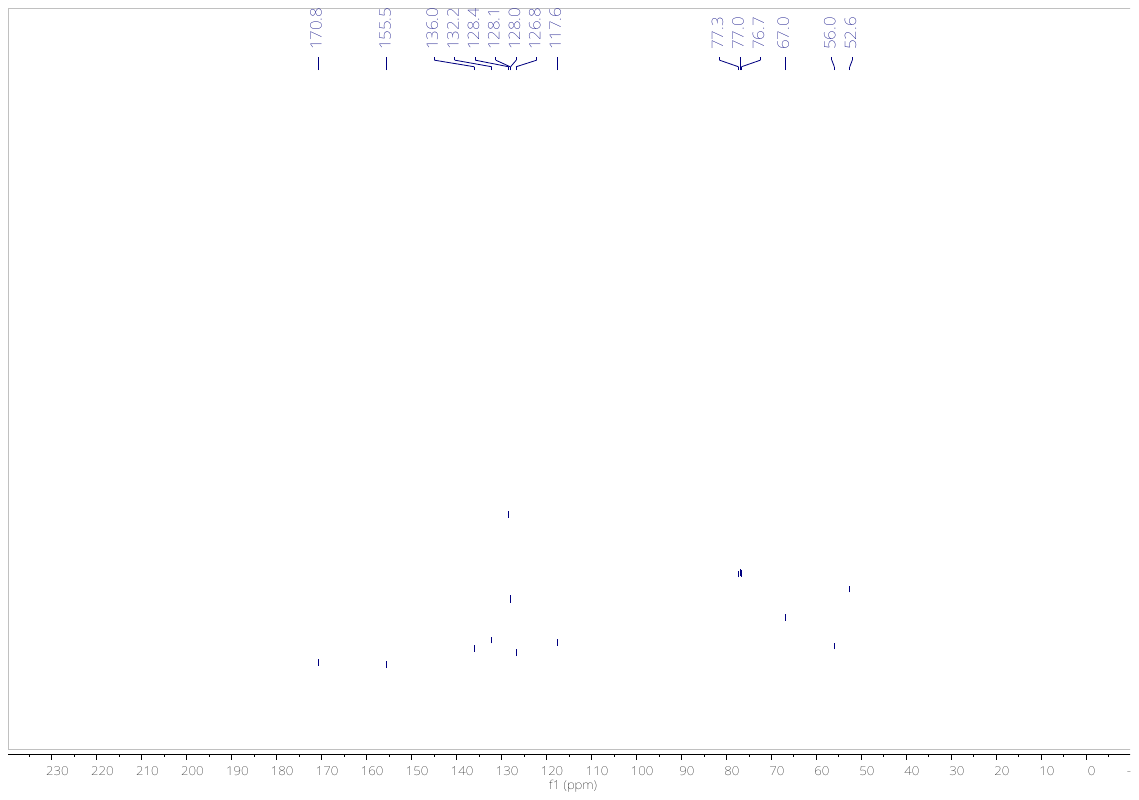

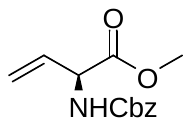

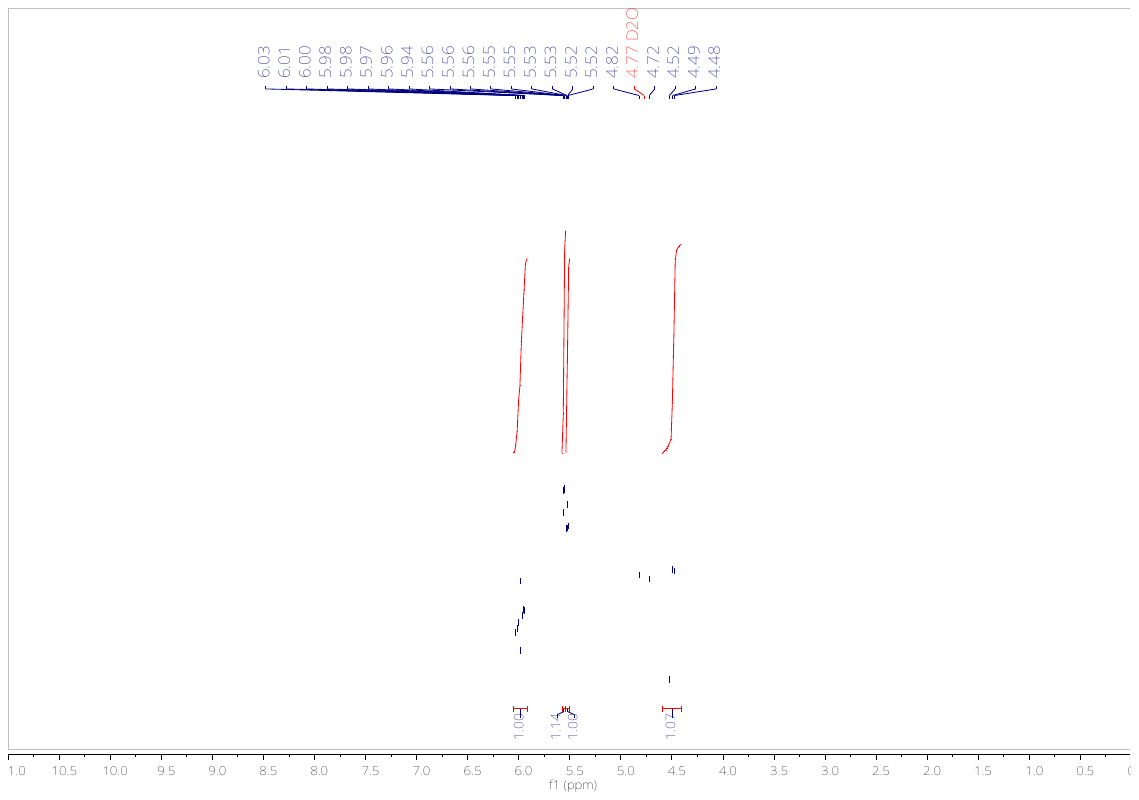

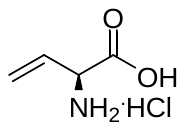

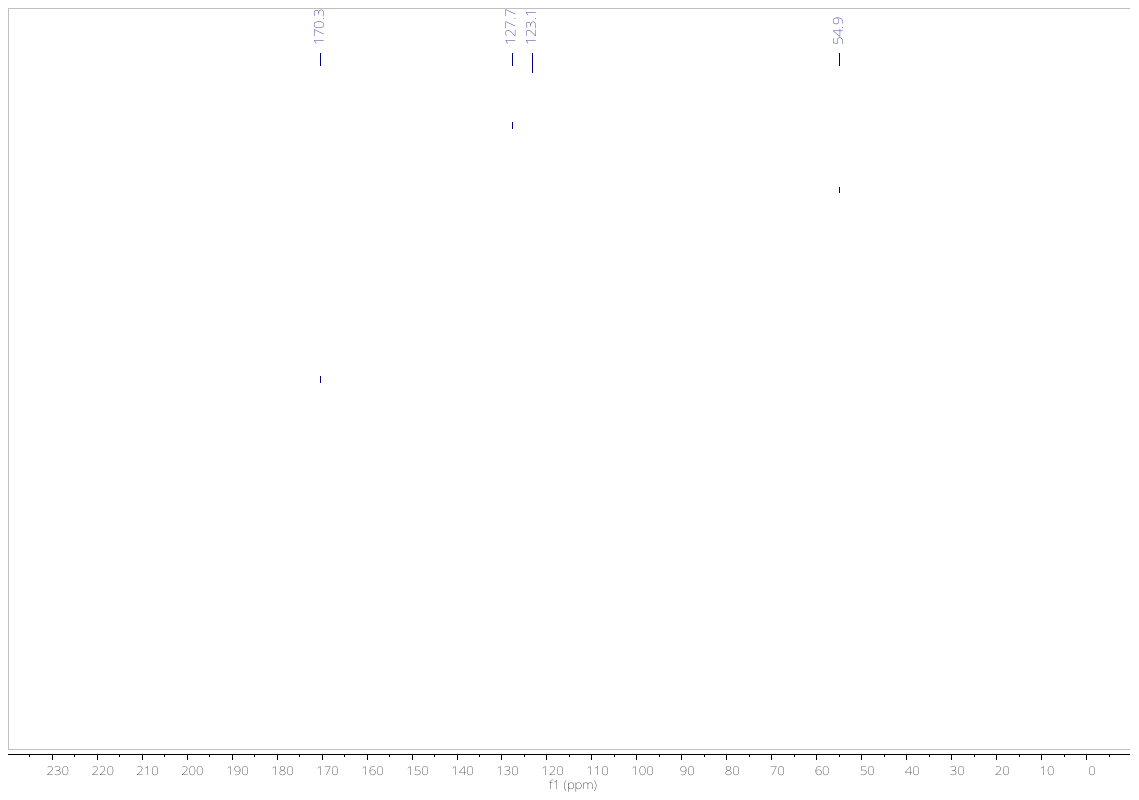

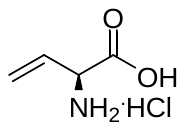

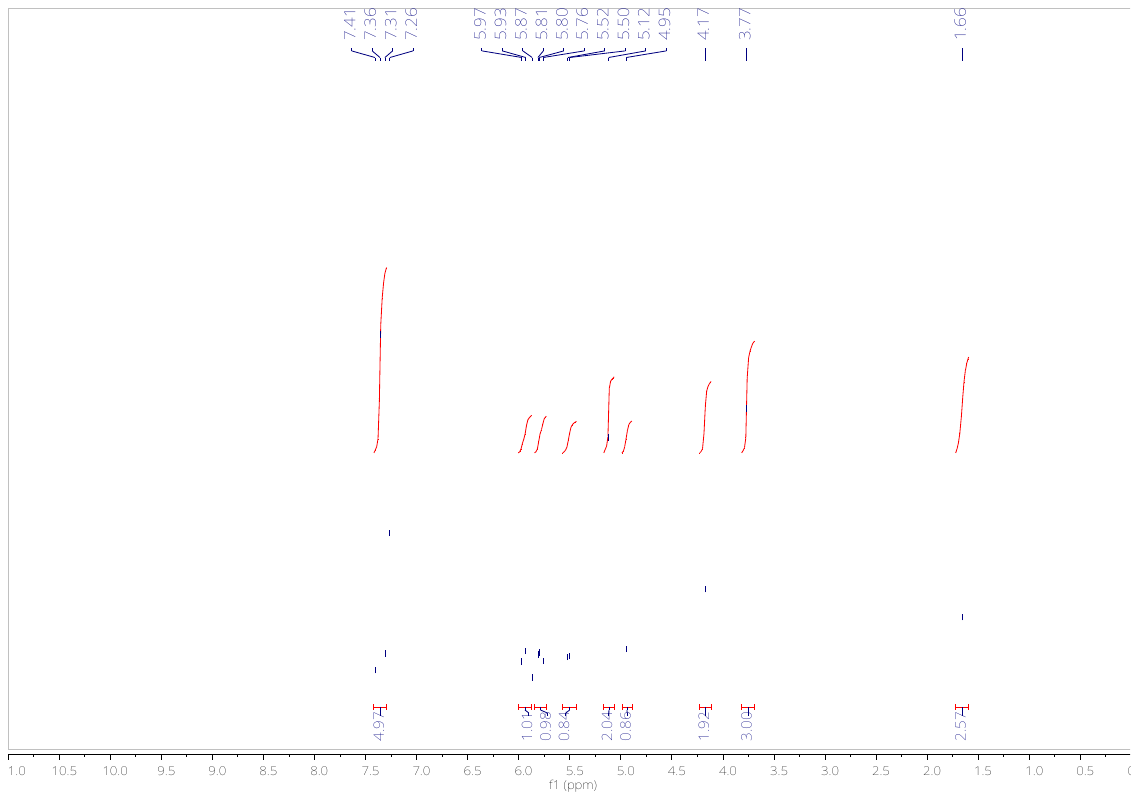

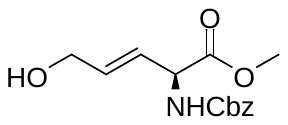

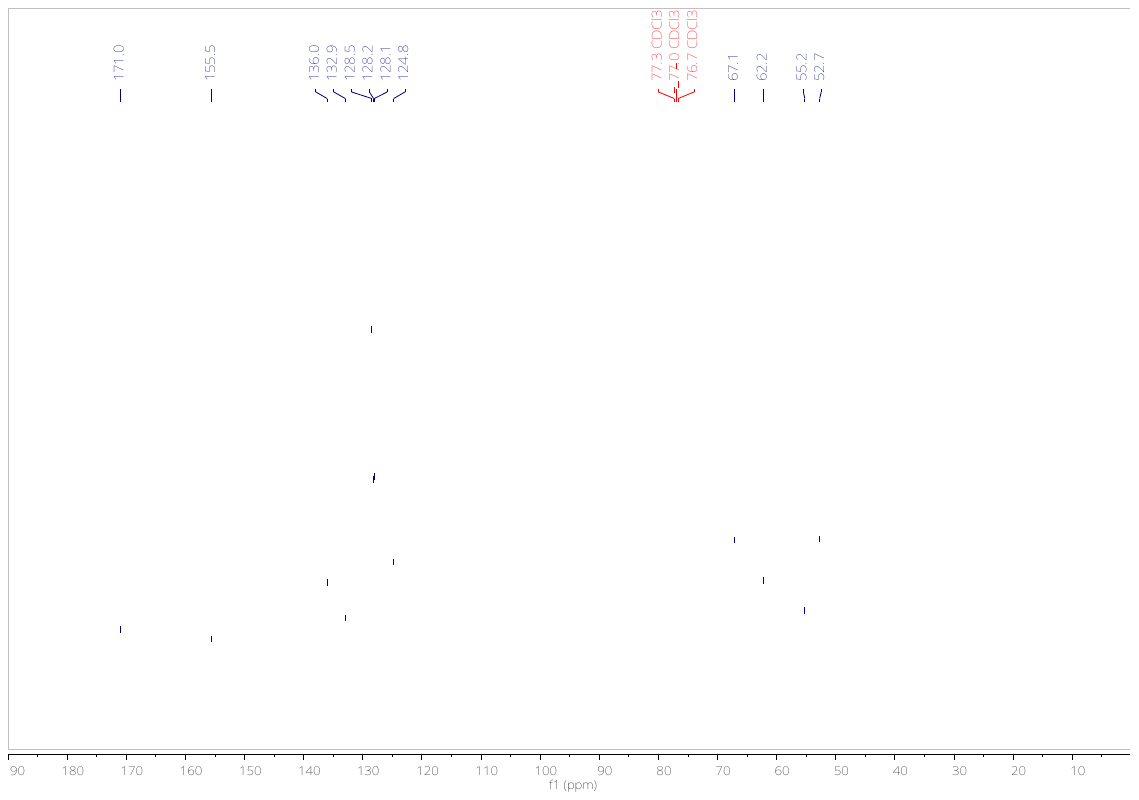

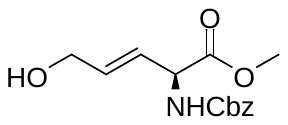

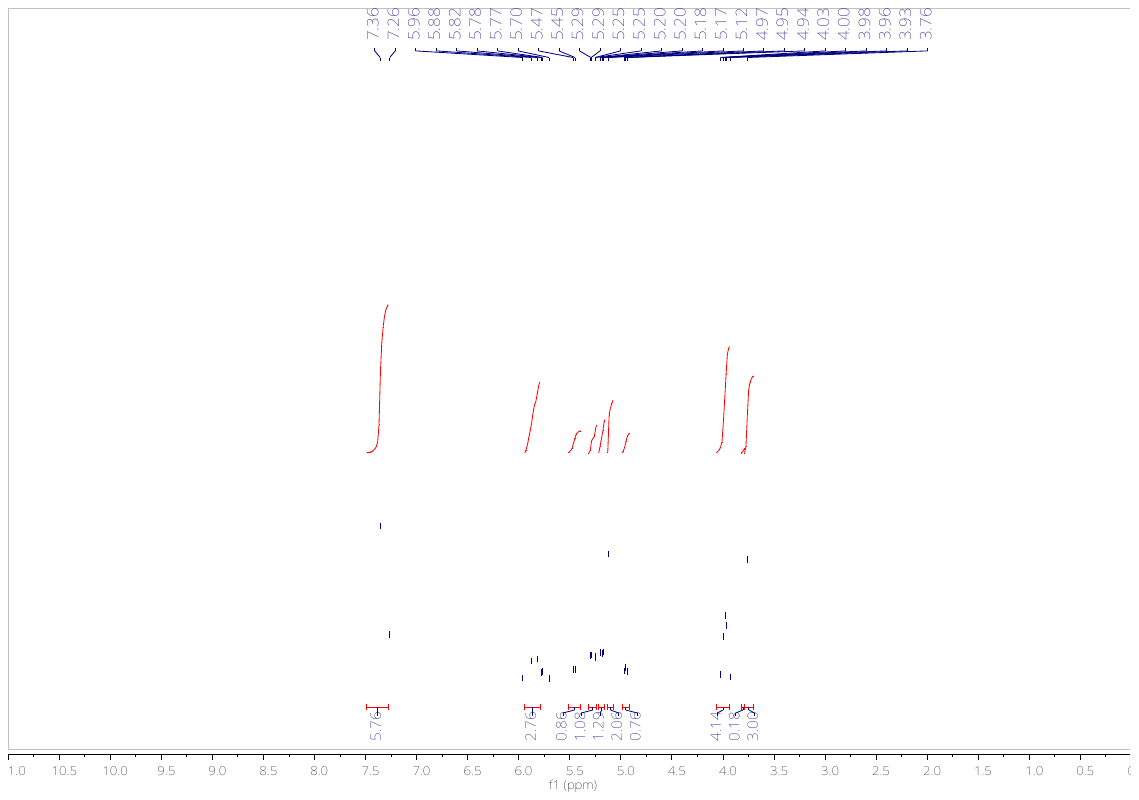

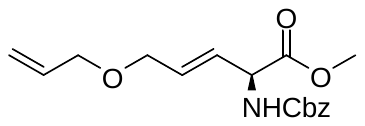

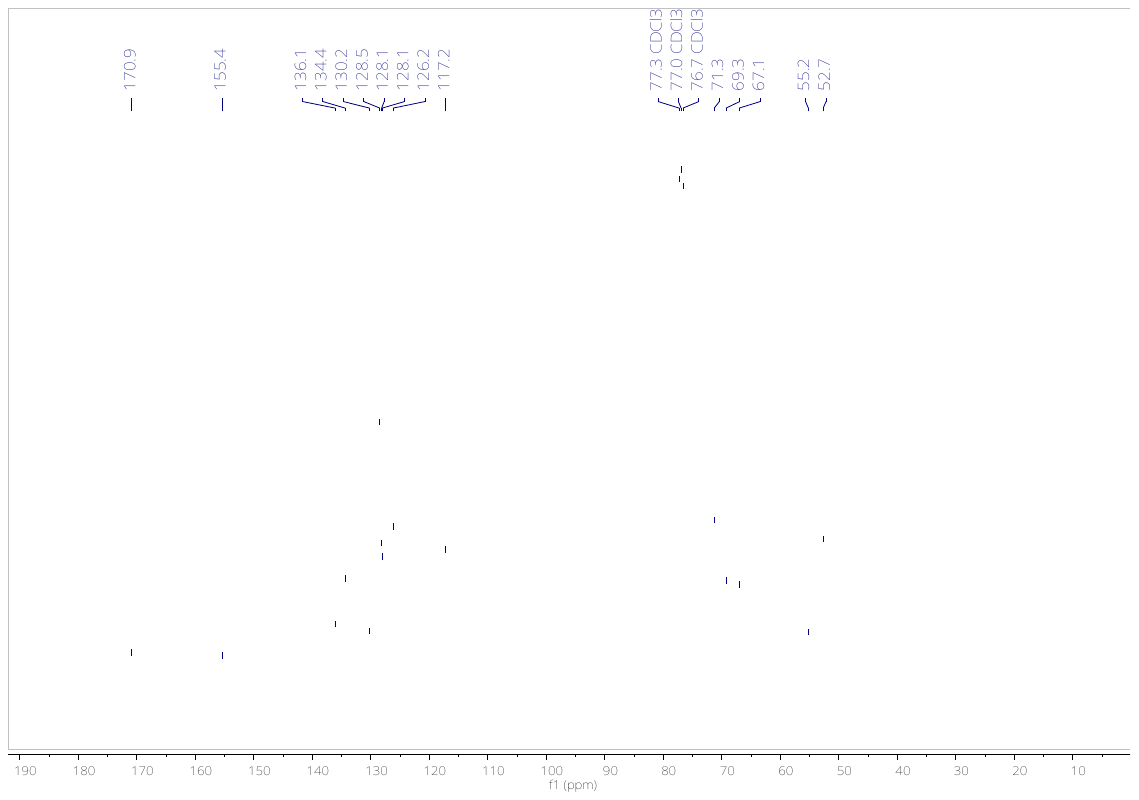

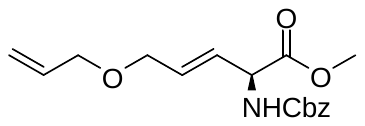

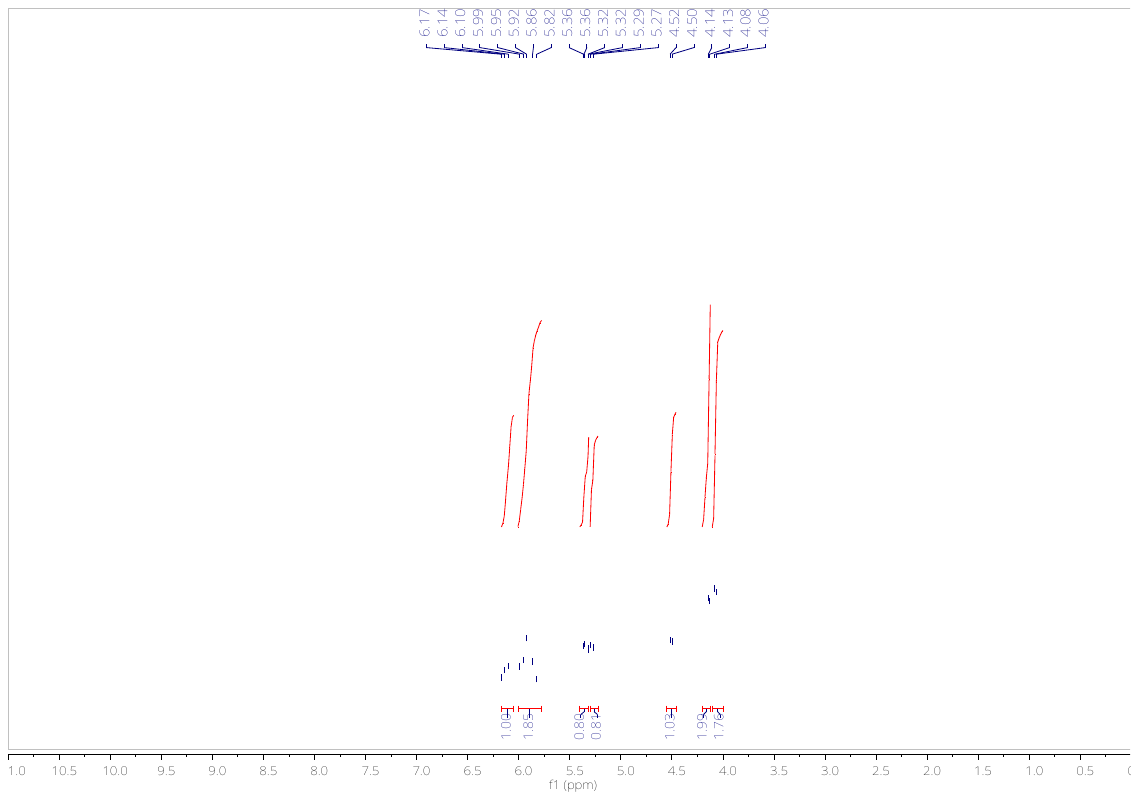

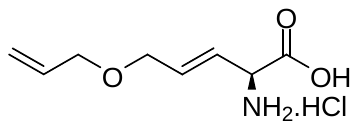

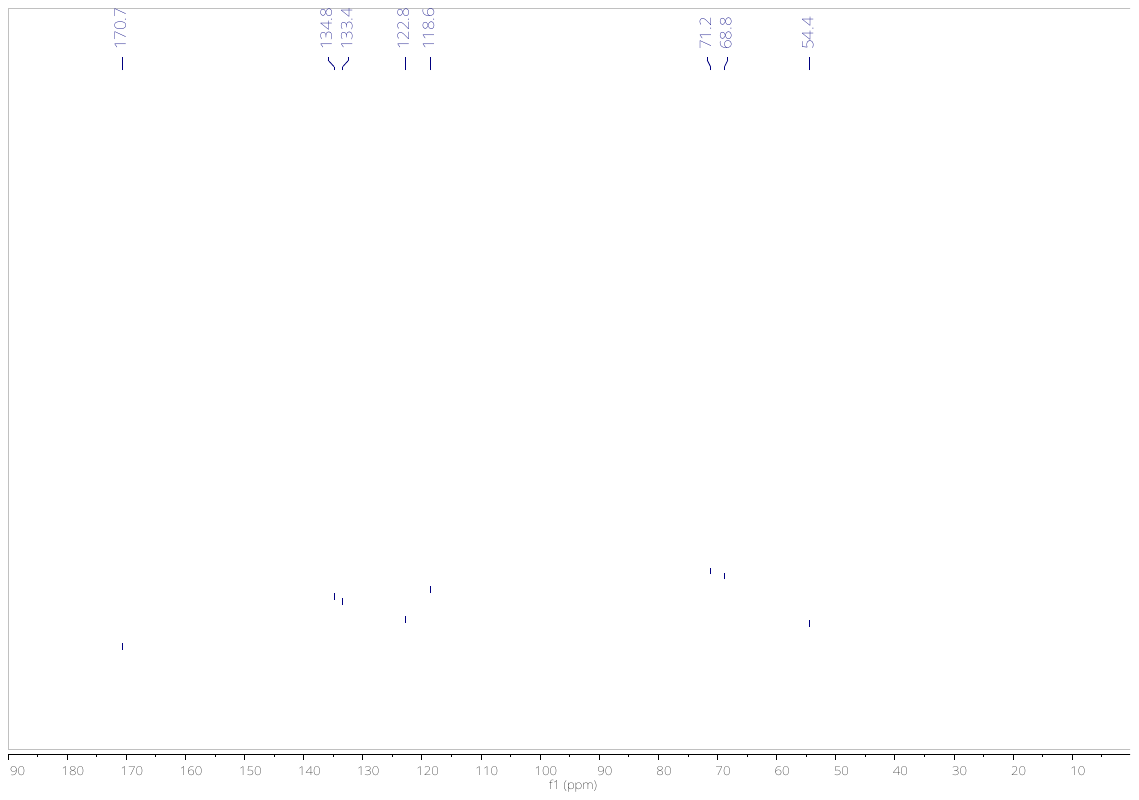

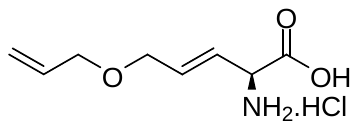
